# Inverse Transformed Encoding Models – a solution to the problem of correlated trial-by-trial parameter estimates in fMRI decoding

**DOI:** 10.1101/610626

**Authors:** Joram Soch, Carsten Allefeld, John-Dylan Haynes

## Abstract

Techniques of multivariate pattern analysis (MVPA) can be used to decode the discrete experimental condition or a continuous modulator variable from measured brain activity during a particular trial. In functional magnetic resonance imaging (fMRI), trial-wise response amplitudes are sometimes estimated from the measured signal using a general linear model (GLM) with one onset regressor for each trial. When using rapid event-related designs with trials closely spaced in time, those estimates are highly variable and serially correlated due to the temporally extended shape of the hemodynamic response function (HRF). Here, we describe inverse transformed encoding modelling (ITEM), a principled approach of accounting for those serial correlations and decoding from the resulting estimates, at low computational cost and with no loss in statistical power. We use simulated data to show that ITEM outperforms the current standard approach in terms of decoding accuracy and analyze empirical data to demonstrate that ITEM is capable of visual reconstruction from fMRI signals.

## 1 Introduction

In functional magnetic resonance imaging (fMRI), data have been traditionally analyzed with *univariate encoding models* (Brodersen et al., 2011b) such as general linear models (GLMs) that construct a relationship between experimental variables and the measured signal in one voxel which allows to statistically test activation differences between experimental conditions (Smith, 2004; Monti, 2011). For some time now, however, data have also been analyzed with *multivariate decoding algorithms* (Brodersen et al., 2011a) such as support vector machines (SVMs) that extract experimental variables from the measured signals in many voxels which allows to reliably decode experimental conditions from brain activation (Haynes and Rees, 2006; Haynes, 2015). These latter techniques are collectively referred to as multivariate pattern analysis (MVPA).

Besides directly decoding from samples extracted from pre-processed fMRI time series, a common approach of MVPA for fMRI consists of calculating *session-wise parameter estimates* and using linear support vector machines (Cox and Savoy, 2003; LaConte et al., 2005) to decode experimental manipulations from multivariate signals in a searchlight moving through the brain (Kriegeskorte et al., 2006; Haynes et al., 2007). However, the same machinery can also be applied to *trial-wise parameter estimates* which can be obtained from post-stimulus time-window averaging (Ress and Heeger, 2003), using a finite impulse response approach (Ress et al., 2000) or via trial-wise response regression (Rissman et al., 2004; Molloy et al., 2018). While the higher number of samples in trial-wise estimates and the lower variance of session-wise estimates both act to increase decoding accuracy and may lead to the same benefit with respect to classification performance, employing trial-wise signals comes closer to the original idea of “decoding” as it allows, for each individual trial, to make a prediction which condition it belongs to.

Trial-wise response amplitudes are most often estimated from the fMRI signal using a GLM with one onset regressor per trial (Rissman et al., 2004) generated by convolution with a *hemodynamic response function* (HRF; Friston et al., 1998; Henson et al., 2001). When using rapid event-related designs with trials closely spaced in time, those estimates are highly variable and serially correlated due to the temporally extended shape of the canonical HRF (Mumford et al., 2012; Turner et al., 2012) which leads to inaccurate parameter estimates and invalid statistical tests (Mumford et al., 2014).

Mumford and colleagues systematically assessed different methods of obtaining trial-wise parameter estimates and found that the so-called “*least squares, separate*” method (LS-S) performed best in terms of the MVPA decoding accuracy among all methods considered (Mumford et al., 2012). The LS-S method obtains each trial’s response via a GLM including a regressor for that trial and another regressor for all other trials (Mumford et al., 2012). Consequently, each trial requires fitting a separate GLM and e.g. calculating activation patterns for 100 trials needs 100 GLMs.

In this work, we introduce a new solution to the problem of correlated trial-by-trial parameter estimates, termed *inverse transformed encoding modelling* (ITEM). Instead of modifying the way how trial-wise response amplitudes are estimated, this solution considers the actual distribution of the trial-wise parameter estimates, as implied by the trial-wise design matrix that is used to generate them. In this way, correlations are not artificially reduced, but naturally accounted for in the subsequent decoding analysis. Importantly, ITEM does not require fitting a separate GLM for each trial, thus extremely lowering the computational cost^1^ of trial-wise MVPA for fMRI.

The structure of this paper is as follows. First, we will outline the theoretical framework underlying ITEM-based analyses (see Section 2). Practitioners not interested in the mathematical details can read a brief summary of the methodology (see Section 2.1). Second, we will perform a simulation study on classification from fMRI data and demonstrate that ITEMs are as powerful as LS-S or, in certain critical situations, even more powerful (see Section 3). Third, we will describe an empirical application in which ITEMs are used for reconstruction of massively parallel visual information in an extremely rapid event-related design (see Section 4). Finally, we will discuss our results (see Section 5).

## 2 Theory

In this section, we introduce the mathematical details of inverse transformed encoding models (ITEMs). Non-technical readers are recommended to read a brief summary of the methodology (see Section 2.1) and then directly proceed to the simulation study (see Section 3) or the empirical validation (see Section 4).

### 2.1 Brief summary of the methodology

In univariate fMRI data analysis, general linear models (GLMs) are commonly used to estimate activation patterns associated with experimental conditions (see Section 2.2). In multivariate fMRI data analysis, a trial-wise GLM is sometimes used to obtain trial-wise response amplitudes on which decoding analyses are then performed (see Section 2.3). We use a mapping between the trial-wise and the standard GLM (see Section 2.4) and derive the full trial-by-trial correlation structure (see Section 2.5) which gives rise to a new model operating on the trial-wise parameter estimates themselves (see Section 2.6). Extending this model to multivariate signals (see Section 2.7) and inverting its explanatory direction (see Section 2.8) allows to classify discrete experimental conditions or reconstruct continuous parametric modulators (see Section 2.9) while at the same time accounting for trial-to-trial correlations due to the slow hemodynamic response.

### 2.2 The standard general linear model

In functional magnetic resonance imaging (fMRI) data analysis, it is common to use *general linear models* (GLMs) for statistical inference (Friston et al., 1994; Friston, 1995; Monti, 2011; Carp, 2012). In a GLM, a single voxel’s fMRI data (*y*) are modelled as a linear combination (*β*) of experimental factors and potential confounds (*X*), where errors (*ε*) are assumed to be normally distributed around zero and to have a known covariance structure (*V*), but unknown variance factor (*σ*^2^):

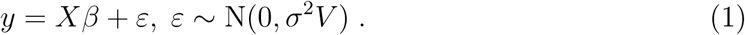

In this equation, *y* is the *n* × 1 measured signal, *X* is the *n* × *p* design matrix, *β* is a *p* × 1 vector of regression coefficients, *ε* is an *n* × 1 vector of errors or noise, *σ*^2^ is the variance of these errors and *V* is an *n* × *n* temporal correlation matrix where *n* is the number of data points and *p* is the number of regressors (see Table 1).

**Table 1.** Mathematical notation. Abbreviations: GLM = general linear model; TEM = transformation encoding model. Categories: data = variable quantities measured during the experiment; model = fixed quantities specified by the researcher; parameter = model parameters that have unknown true values; estimate = estimated values of those model parameters; noise = parts of the data that are assumed to be random. Note that when *r >* 0, dimensions *t* and *p* become (*t* + *r*) and (*p* + *r*), respectively.

| Symbol | Dimension | Category | Description |
| --- | --- | --- | --- |
| dimensions of data, experiment and model |  |  |  |
| $n$ | scalar | – | number of fMRI scans |
| $t$ | scalar | – | number of trials in the experiment |
| $p$ | scalar | – | number of experimental conditions |
| $r$ | scalar | – | number of nuisance regressors |
| $v$ | scalar | – | number of voxels (e.g. in ROI) |
| standard GLM: $y = X\beta + \varepsilon$, $\varepsilon \sim N(0, \sigma^2 V)$ [Section 2.2] | | | |
| $y$ | $n \times 1$ | data | a single voxel's fMRI signal |
| $X$ | $n \times p$ | model | design matrix of the standard GLM |
| $\beta$ | $p \times 1$ | parameter | condition-wise response amplitudes |
| $\hat{\beta}$ | $p \times 1$ | estimate | condition-wise parameter estimates |
| $\varepsilon$ | $n \times 1$ | noise | errors/noise of the standard GLM |
| $\sigma^2$ | scalar | parameter | noise variance of the standard GLM |
| $V$ | $n \times n$ | model | scan-by-scan covariance matrix |
| trial-wise GLM: $y = X_t\gamma + \varepsilon_t$, $\varepsilon_t \sim N(0, \sigma_t^2 V)$ [Section 2.3] | | | |
| $X_t$ | $n \times t$ | model | design matrix of the trial-wise GLM |
| $\gamma$ | $t \times 1$ | parameter | trial-wise response amplitudes |
| $\hat{\gamma}$ | $t \times 1$ | estimate | trial-wise parameter estimates |
| $\varepsilon_t$ | $n \times 1$ | noise | errors/noise of the trial-wise GLM |
| $\sigma_t^2$ | scalar | parameter | noise variance of the trial-wise GLM |
| univariate TEM: $\hat{\gamma} = T\beta + \eta$, $\eta \sim N(0, \sigma^2 U)$ [Section 2.6] | | | |
| $\hat{\gamma}$ | $t \times 1$ | estimate | estimated trial-wise response amplitudes |
| $T$ | $t \times p$ | model | design matrix of the univariate TEM |
| $\eta$ | $t \times 1$ | noise | noise vector of the univariate TEM |
| $U$ | $t \times t$ | model | trial-by-trial covariance matrix |
| multivariate TEM: $\hat{\Gamma} = TB + H$, $H \sim MN(0, U, \Sigma_y)$ [Section 2.7] | | | |
| $\hat{\Gamma}$ | $t \times v$ | estimate | trial- and voxel-wise parameter estimates |
| $B$ | $p \times v$ | parameter | activation pattern of the multivariate TEM |
| $H$ | $t \times v$ | noise | noise matrix of the multivariate TEM |
| $\Sigma_y$ | $v \times v$ | parameter | voxel-by-voxel covariance matrix |
| inverse TEM: $T = \hat{\Gamma}W + N$, $N \sim MN(0, U, \Sigma_x)$ [Section 2.8] | | | |
| $W$ | $v \times p$ | parameter | extraction filter of the inverse TEM |
| $\hat{W}$ | $v \times p$ | estimate | estimated extraction filter = weight matrix |
| $\hat{W}_{-j}$ | $v \times p$ | estimate | weight matrix, estimated from training set |
| $\hat{T}_j$ | $t \times p$ | estimate | design variables, decoded in test set |
| $N$ | $t \times p$ | noise | noise matrix of the inverse TEM |
| $\Sigma_x$ | $p \times p$ | parameter | condition-by-condition covariance matrix |

The design matrix *X* usually consists of stimulus functions representing experimental conditions which are convolved with a hemodynamic response function (HRF; Friston et al., 1998; Henson et al., 2001) and a set of nuisance regressors not based on HRF convolution such as movement parameters. The covariance structure *V* is, at least in Statistical Parametric Mapping (SPM; Friston et al., 2007), obtained from fitting an AR(1) model to fMRI signals from all active voxels in the brain (Friston et al., 2002a; Friston et al., 2002b), such that it is considered as known for each individual voxel. Given known *X* and *V* as well as measured *y*, maximum likelihood estimates for the regression coefficients can be obtained via weighted least squares (WLS) as

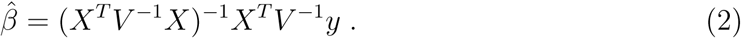

Based on estimated model parameters 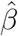, classical statistical inference can be performed by defining *t*- or *F*-contrasts, calculating the respective *t*- and *F*-statistics and comparing them to the *t*- or *F*-distribution under the respective null hypothesis (Ashburner et al., 2003, ch. 8; Friston et al., 2007, ch. 9).

### 2.3 The trial-wise general linear model

The standard GLM for fMRI makes the assumption that all trials within one condition, i.e. all events in one column of the design matrix *X*, elicit the same response in the measured signal *y*. If we wish to relax this assumption or if we want to analyze trial-wise responses separately, we can specify a *trial-wise general linear model* :

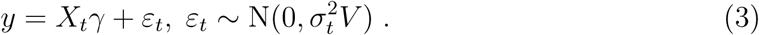

In this equation, *X*_*t*_ is an *n* × *t* trial-wise design matrix, *γ* is a *t* × 1 vector of trial-wise response amplitudes, *ε*_*t*_ is an *n* × 1 vector of errors and 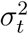 is the variance of these errors where *t* is the number of trials in the experiment (see Table 1).

More precisely, *X*_*t*_ is a matrix with one column for each trial and each column consists of one single event, convolved with a hemodynamic response function (see Figure 1A). This allows to obtain trial-wise parameter estimates:

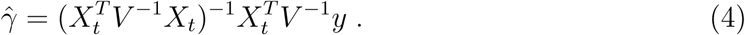

**Figure 1.**
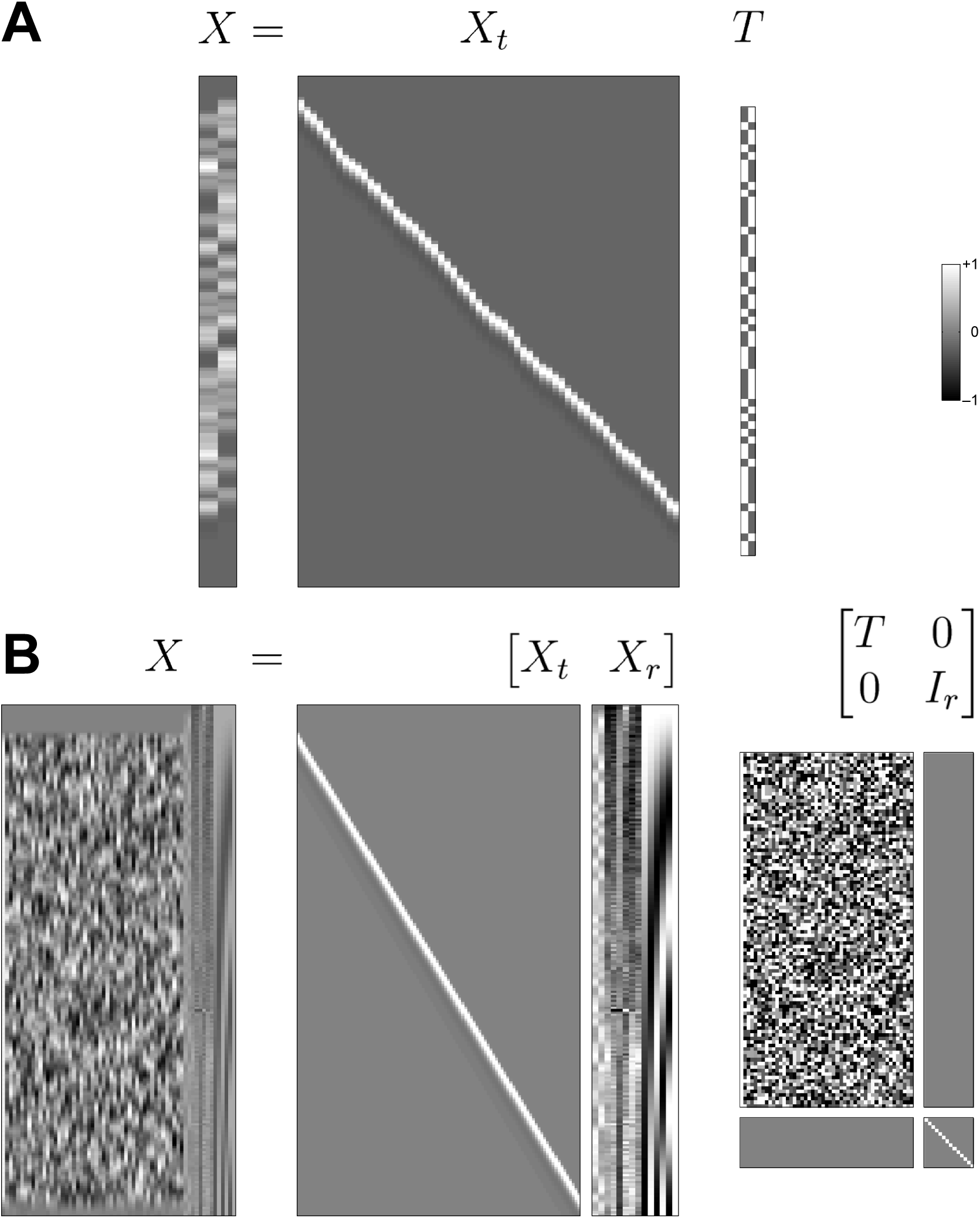
The transformation matrix. This figure illustrates how the *T* matrix maps from the trial-wise design matrix *X*_*t*_ to the standard design matrix *X*. Generally, *T* has as many rows as *X*_*t*_ has columns (the number of trials) and as many columns as *X* (the number of conditions). (A) In a very simple case (from our simulation example, see Section 3), *T* is just a binary indicator matrix that collects individual trials from *X*_*t*_ into experimental conditions in *X*. (B) In a more complicated case (from our empirical validation, see Section 4), *T* also has columns with continuous values to emulate parametric modulators in *X* and an identity matrix at the bottom right to append nuisance regressors to *X*. For a detailed description of the regressors in *X*, see Section 4.2. Note that pixel sizes are not identical across matrices, but optimized for visibility purposes.

Those values form the basis for the commonly known “least squares, all” (LS-A) method (Mumford et al., 2012) which operates on these raw parameter estimates. They are also used in the ITEM approach, but with the crucial difference that their trial-by-trial co-variance is being accounted for in ITEM (see next sections).

### 2.4 The transformation matrix *T*

The standard GLM and the trial-wise GLM are two different encoding models for univariate, i.e. single-voxel fMRI data. The standard GLM allows to estimate condition-specific effects and contrast them for statistical inference (Friston et al., 1994) whereas the trial-wise GLM allows to estimate trial-wise response amplitudes from the BOLD signal (Rissman et al., 2004).

Typically, the *n* × *p* design matrix *X* has a scan-by-regressor structure where each row corresponds to one fMRI scan and each column corresponds to one experimental condition (see Figure 1A), i.e. stimulus onsets and durations, convolved with the canonical HRF. In contrast, the *n* × *t* design matrix *X*_*t*_ has a scan-by-trial structure so that each column corresponds to one event (see Figure 1A) and basically is an onset regressor with a single HRF at the time of the corresponding trial. The core idea of this contribution is to connect these matrices via the relation

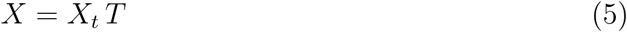

where the *t* × *p transformation matrix T* is defined such that it converts trial-wise HRFs into condition regressors (see Figure 1). In the case of a purely categorical design, *T* will simply be a binary indicator matrix where *t*_*ij*_ = 1 indicates that the *i*-th trial belongs to the *j*-th condition (see Figure 1A). If there is a parametric modulator in the design matrix, *T* will have a corresponding column with the modulator values belonging to this regressor (see Figure 1B).

Usually, the design matrix also includes nuisance regressors *X*_*r*_, e.g. events of no interest, movement parameters, filter regressors or the implicit baseline (see Figure 1B). These nuisance regressors prohibit a trial-to-scan mapping since they are not based on trial-wise modulation. In this case, to preserve equation (5), these regressors are simply appended to *X*_*t*_ and *T* takes on a block-diagonal structure as

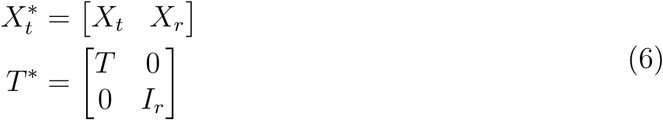

where 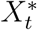 is the *n* × (*t* +*r*) “augmented” trial-wise design matrix, *T* ^*^ is the (*t* +*r*) × (*p* +*r*) “augmented” transformation matrix (see Figure 1B) and *r* is the number of nuisance regressors. In what follows, when we use the symbols *X*_*t*_ and *T* as well as *t* and *p*, we almost always refer to the augmented quantities 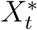 and *T* ^*^ as well as (*t* + *r*) and (*p* + *r*).

### 2.5 The uncorrelation matrix *U*

Given that trial-wise parameter estimates 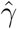 – representing BOLD signal response amplitudes during individual trials – have been estimated from the data via (4), there will be a certain covariance between them due to the fact that trial-wise HRF regressors overlap and are thus temporally correlated with each other. It can be shown that this covariance is a function of the trial-wise design matrix *X*_*t*_ (see Appendix A, Theorem 1):

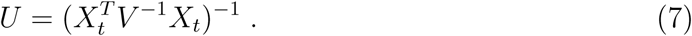

We refer to this matrix as the *uncorrelation matrix*, because it allows to decorrelate trial-wise response amplitudes^2^ when their HRFs are overlapping in time. Using this covariance matrix that directly derives from the trial-wise design matrix, the correlation between adjacent trials imposed by temporally close HRFs can be easily accounted for in a second model on the trial-wise parameter estimates (see next section).

Notably, the *U* matrix does not only capture correlations between trial-wise parameter estimates alone (see Figure 2A), but also accounts for possible correlations between trial-wise HRFs and nuisance variables such as filter regressors (see Figure 2B). This suggests not to regress nuisance variables beforehand, but instead to include all processes of interest and of no interest into the model at once (see Figure 1B).

**Figure 2.**
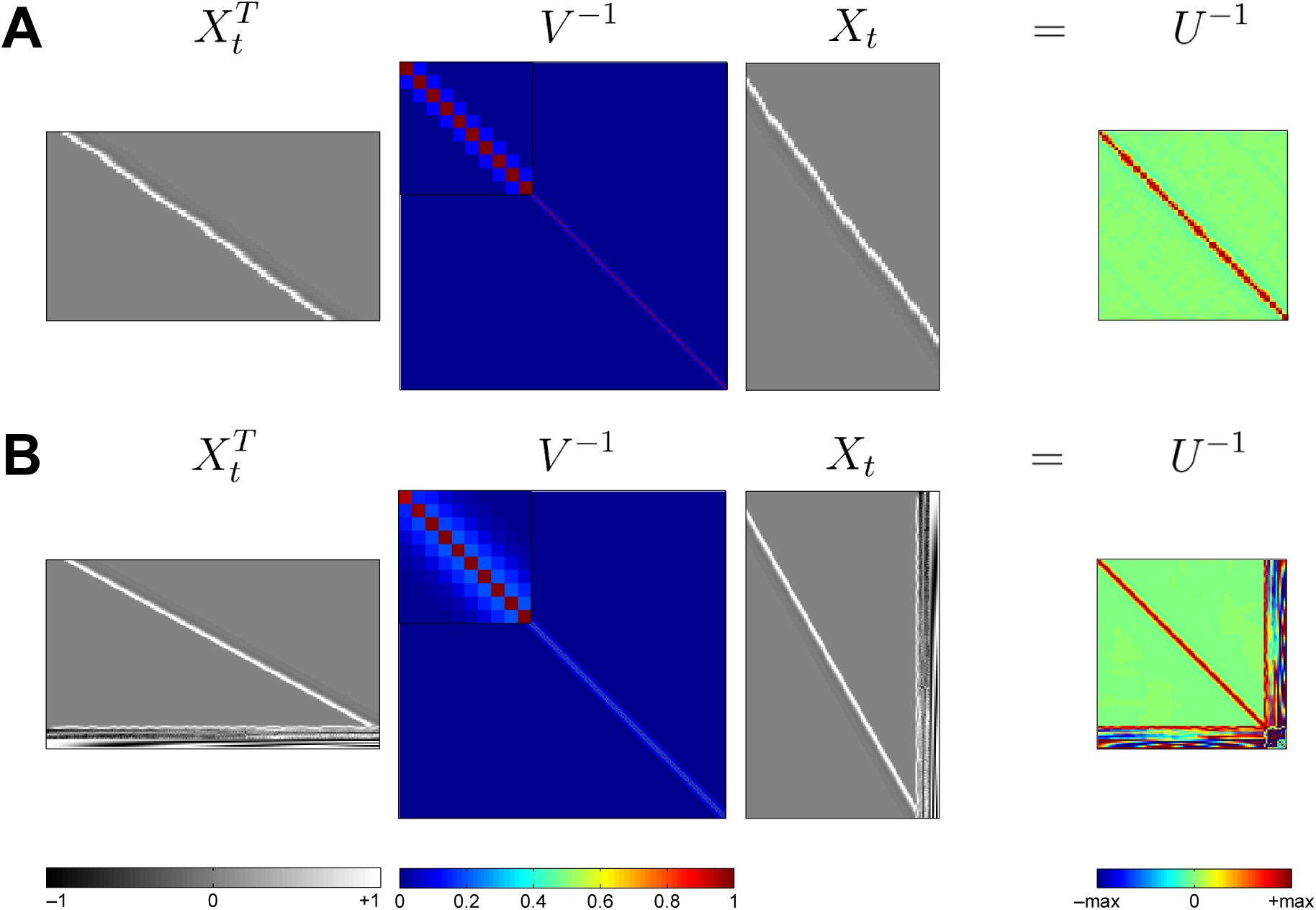
The uncorrelation matrix. This figure illustrates how the *U* matrix derives from the trial-wise design matrix *X*_*t*_ and scan-to-scan covariance matrix *V*. Generally, the inverse of *U* is a product of *X*_*t*_ with itself, weighted by the inverse of *V*. (A) In a very simple case (from our simulation example, see Section 3), *U* only encodes correlations between adjacent trials. The closer two trials are to each other in time, the stronger are their HRFs correlated, illustrated by red entries in *U*^−1^ for short inter-stimulus-intervals in *X*_*t*_. (B) In a more complicated case (from our empirical validation, see Section 4), the augmented 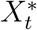 also includes nuisance regressors, such that *U* not only encodes trial-by-trial correlations (upper left portion), but also the shared variation between trial-wise HRFs and regressors of no interest (rightmost columns and lowermost rows). The upper-left insets in the plots of *V*^−1^ correspond to the (inverse) covariance pattern between ten consecutive scans.

Note that the present derivation is based on assuming constant trial-wise response amplitudes within experimental conditions^3^. If this assumption is to be relaxed, the covariance of the trial-wise parameter estimates becomes more complicated (see Appendix B) and restricted maximum likelihood (ReML) estimation is required. We have used this ReML extension in the simulation study of this paper (see Section 3), but have found no evidence of improvement in empirical analyses (see Section 4), and therefore only present it as a possible extension of our framework (see Appendix B).

### 2.6 The transformed encoding model

We now assume that 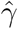, *T* and *U* are given from equations (4), (5) and (7), respectively. Together, the transformation matrix *T* and the uncorrelation matrix *U* can be used to define a new linear model on the trial-wise parameter estimates 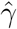, by specifying their distribution resulting from the estimation (see Appendix A, Theorem 1):

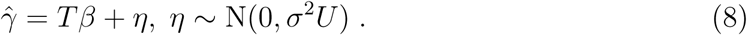

We refer to this trial-level GLM as a *transformed encoding model* (TEM), because it operates on a transformed version of the data, namely the trial-wise response amplitudes 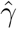, and uses the transformation matrix *T* as its design matrix.

Other than in *X* or *X*_*t*_, where information about trials is only indirectly contained (because convolved), it is directly accessible via the rows of *T* which makes the model suitable for trial-wise decoding analyses. When adopting this model, condition-wise parameter estimates 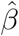 can be derived from the trial-wise parameter estimates 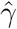 via

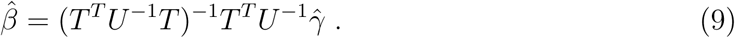

and it can be shown (see Appendix A, Theorem 2) that they are identical to the condition-wise parameter estimates of the standard GLM given by (2).

### 2.7 The multivariate transformed encoding model

Given that trial-wise response amplitudes 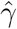 have been estimated in a number of voxels, e.g. a searchlight (SL) or a (functional or anatomical) region of interest (ROI), we can turn the univariate GLM (8) into a multivariate GLM (Allefeld and Haynes, 2014), the *multivariate transformed encoding model* (MTEM):

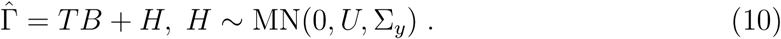

In this equation, 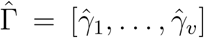 is a *t* × *v* matrix of horizontally concatenated trial responses over voxels, *B* and *H* are the corresponding extensions of *β* and *η*, MN indicates a matrix-normal distribution and σ_*y*_ is the *v* × *v* unknown spatial covariance matrix where *v* is the number of voxels currently analyzed.

With the transition from the univariate to the multivariate TEM, we are now able to describe multi-voxel activation patterns rather than single-voxel response amplitudes. Importantly, while the voxel-to-voxel covariance σ_*y*_ changes depending on the set of voxels considered and actually allows for the multivariate encoding exploited in decoding analyses, the trial-to-trial covariance remains the same, namely *U*, because it only depends on the trial-wise design matrix *X*_*t*_ and the scan-to-scan covariance matrix *V* used to generate them, as indicated by equations (4) and (7). Because *V* is usually estimated from whole-brain data (see Section 2.2) and considered identical across voxels, *U* will also be the same for each set of voxels considered.

### 2.8 The inverse transformed encoding model

In principle, our investigation could stop here and data analysis could proceed by statistically inferring single-voxel activation differences using the univariate TEM (see Section 2.6) or multi-voxel pattern differences using the multivariate TEM (see Section 2.7). This would only require to transform measured data *Y* into trial-wise parameter estimates 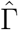 via (4), to incorporate the trial-to-trial covariance *U* calculated from (7) and could operate in the standard frameworks for the univariate GLM (Friston et al., 1994) and the multivariate GLM (Allefeld and Haynes, 2014).

However, our goal here is not statistical inference, i.e. describing the trial-wise response amplitudes 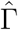 in terms of the experimental design *T*, but decoding analysis, i.e. recovering the experimental design from trial-wise response amplitudes. Therefore, we define an extraction filter *W* as the inverse of the activation pattern *B*. Then, it can be shown (see Appendix C, Theorem 4) that the forward GLM (10) implies the following backward GLM, the *inverse transformed encoding model* (ITEM):

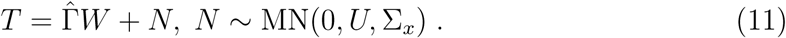

In this equation, the known transformation matrix *T* occurs as the “data” matrix, the estimated response amplitudes 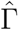 become the “design” matrix^4^, *W* appears as the unknown *v* × *p* “weight” matrix, *N* is a *t* ×*p* “noise” matrix and σ_*x*_ is the unknown *p* × *p* covariance matrix across experimental design variables.

Based on this model, an estimate of the extraction filter, also called a “weight matrix”, can be obtained using weighted least squares which is the best linear unbiased estimator (BLUE) in this situation (see Appendix C, Theorem 5):

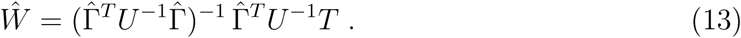

In the present work, to assess decoding accuracy, ITEMs will be estimated by cross-validation (CV) across fMRI recording sessions. More precisely, we will perform leave-one-session-out cross-validation across *S* sessions. In each CV fold *j* = 1, …, *S*, a weight matrix *Ŵ*_¬*j*_ is calculated from all except the *j*-th session^5^:

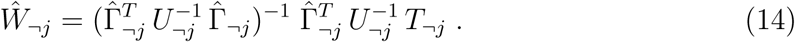

Then, this weight matrix is used to obtain decoded design variables 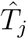 via simple out-of-sample prediction in the left-out session *j*:

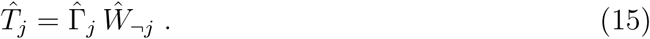

### 2.9 Decoding by classification or reconstruction

In the ITEM framework, *decoding* is generally understood as recovering an independent variable and two types of decoding analysis are possible: (i) *classification*, i.e. decoding discrete categories, e.g. experimental conditions; and (ii) *reconstruction*, i.e. decoding continuous variables, e.g. parametric modulators.

In cases of *classification*, we measure decoding accuracy based on *proportion correct*. First, a *p* × *q* contrast matrix *C* is defined where *q* is the number of classes to discriminate and *c*_*ij*_ = 1 indicates that the *i*-th regressor in *T* belongs to the *j*-th class.^6^ The design variables to be decoded and plugged into (14) are then given by

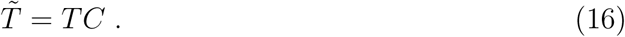

When predicted variables have been calculated from (15), decoding accuracy is determined as the proportion of trials in which the class with the largest decoded value is identical with the class that was actually present, i.e.^7^

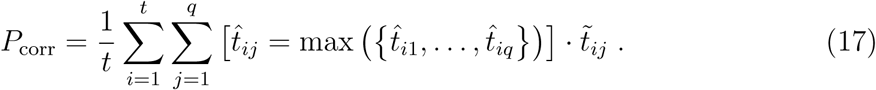

In cases of *reconstruction*, we measure decoding accuracy via *correlation coefficients*. First, a *p* × *q* contrast matrix *C* is defined where *q* is the number of variables to reconstruct and *c*_*ij*_ = 1 indicates that the *i*-th regressor should be evaluated.^8^ Then, the design variables to be decoded are again given by equation (16).

When predicted variables have been calculated from (15), decoding accuracy is determined as the Pearson correlation between original regressor and reconstructed regressor for each variable of interest, i.e.

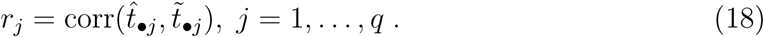

Note that, when *T* has been reconstructed via 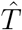, any desired measure of decoding accuracy may be applied. For example, if parametric regressors represent basis functions over stimulus space (e.g. Brouwer and Heeger, 2009), it may be more informative to recover the stimulus by combining information across reconstructed basis functions within trials (e.g. Sprague et al., 2016, suppl. eq. 5) rather than calculating the reconstruction accuracy of each basis function across trials.

An overview of our modelling logic is given in Figure 3. In this demonstration example, we generated trial-wise response amplitudes 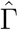 using equation (10) based on ground truth values for *T, B, U* and σ_*y*_ (see Figure 3A). Then, estimation of *Ŵ* using equation (13) and predicition of 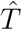 using equation (15) without cross-validation gave rise to a within-sample reconstruction of *T* (see Figure 3B). Finally, we calculated *BŴ* to validate assumptions made in our derivations (see Figure 3C).

**Figure 3.**
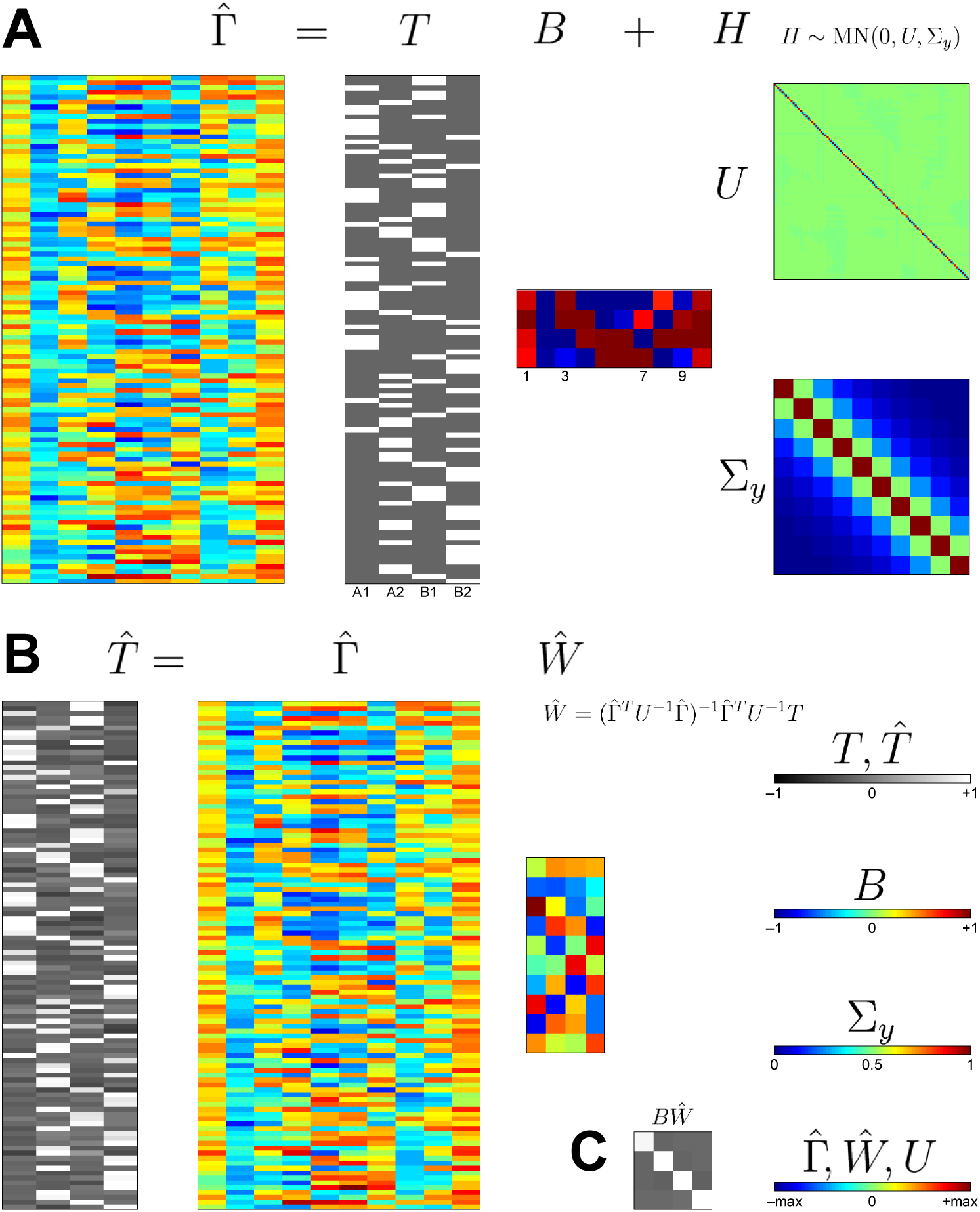
A hypothetical reconstruction. This figure summarizes our approach and illustrates trial-wise reconstruction of experimental design information. (A) Using the multivariate transformed encoding model (see Section 2.7), the trial-wise parameter estimates 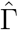 are assumed to be a linear combination of experimental conditions *T*, weighted by voxel-wise regression coefficients *B*, and a noise matrix *H* with trial-by-trial correlations *U* and voxel-to-voxel covariance σ_*y*_. For this example, *T* and *U* were obtained from an SPM template data set on face repetition priming (Henson et al., 2002) while *B* and σ_*y*_ were chosen as ground truth. *B* was set such that voxels exhibit an overall effect (e.g. voxel 1), main effects (voxels 3 & 7) or an interaction effect (voxel 9) in the 2 × 2 design. (B) Using the inverse transformed encoding model (see Section 2.8), a reconstruction 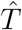 is obtained by multiplying the trial-wise response amplitudes 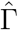 with a weight matrix *Ŵ*, estimated from the inverse model. As one can see when comparing 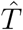 with *T*, experimental conditions can be read out from the reconstruction (see Section 2.9). (C) The product of activation pattern and estimated weight matrix, *BŴ*, approximates the identity matrix – confirming an assumption made when deriving the inverse model (see Appendix C).

## 3 Simulation

### 3.1 Methods

For simulation validation, we repeat and adapt a simulation reported earlier (Mumford et al., 2012) that was designed to investigate different methods of obtaining trial-wise response amplitudes for multivariate pattern analysis in fMRI. All simulation code is available from GitHub (https://github.com/JoramSoch/ITEM-paper).

In our simulation, we compare three approaches of inference from trial-wise parameter estimates: the naïve approach ignoring trial-by-trial correlations (Mumford: “least squares, all”, LS-A), the state-of-the-art approach found optimal in the previous simulation (Mumford: “least squares, separate”, LS-S) and the approach proposed here, i.e. inverse transformed encoding modelling (ITEM). LS-A entails decoding without accounting for correlation and taking trial-wise parameter estimates from equation (4) “as is”. Within our framework, this is equivalent to setting *U* = *I*_*t*_ instead of taking *U* from equation (7), i.e. assuming no correlation between adjacent trials. LS-S cannot be represented within our approach, because it is based on obtaining trial-wise parameter estimates using a separate design matrix for each trial, including one regressor for this trial and one regressor for all other trials (Mumford et al., 2012).

In the simulation, data were generated as follows: First, trials were randomly sampled from two experimental conditions, A and B. Second, trial-wise response amplitudes *γ* were sampled from normal distributions with expectations *µ*_*A*_ = 5 and *µ*_*B*_ = 3 and variances 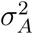 and 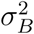 where *σ*_*A*_ = *σ*_*B*_ = 0.5. Third, inter-stimulus-intervals *t*_isi_ were sampled from a uniform distribution U(*t*_min_, *t*_max_) where *t*_min_ ∈ {0, 2, 4} and *t*_max_ = *t*_min_ + 4 seconds. Fourth, the design matrix *X*_*t*_ was generated based on the inter-stimulus-intervals *t*_isi_ and convolution with the canonical hemodynamic response function (cHRF). An exemplary design matrix for the case *t*_isi_ ~ U(0, 4) is given on the left of Figure 1A. Finally, a univariate signal was generated by multiplying the trial-wise design matrix *X*_*t*_ with trial-wise response amplitudes *γ* and adding zero-mean Gaussian noise *ε* with variance *σ*^2^ where *σ* ∈ {0.8, 1.6, 3}. In this way, *N* = 1,000 simulation runs with *S* = 2 sessions (for cross-validation) and *t* = 60 trials (30 per condition) were performed. A detailed description of the simulation and our modifications is given in Appendix E.

After data generation, models were estimated as follows: For LS-A and ITEM, trial-wise parameter estimates 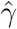 were obtained by equation (4) using design matrix *X*_*t*_ (Mumford: *X*_*S*_). For ITEM, 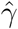 was subjected to an additional restricted maximum likelihood (ReML) analysis, as described in Appendix B, in order to separate the *natural* trial-to-trial variability (coming from 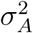 and 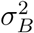) from the *induced* trial-by-trial correlations (coming from *X*_*t*_). For LS-S, 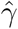 was obtained as described above using trial-specific design matrices *X*_*i*_ where *i* = 1, …, *t* (Mumford: *X*_*T i*_). Afterwards, parameter estimates were subjected to a two-sample t-test in order to assess statistical power (by calculating the proportion of positive results when the alternative hypothesis is true) and a logistic regression in order to assess decoding accuracy (see next section).

### 3.2 Results

The present simulation entails a comparison between activation patterns from two experimental conditions. The classical equivalent to decoding between two conditions is a two-sample t-test. For LS-A and LS-S, trial-wise parameter estimates 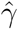 are simply grouped into two vectors which are t-tested against each other. For ITEM, in order to account for correlations in trial-wise parameter estimates 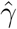, the transformed encoding model given by (8) is estimated using (9) with *T* and *U* given by Figures 1A and 2A, respectively. Then, standard contrast-based inference, as implemented in SPM, is performed (Ashburner et al., 2003, ch. 8; Friston et al., 2007, ch. 9).

Each procedure leads to one *p*-value per simulation and the null hypothesis *H*_0_ : *µ*_*A*_ = *µ*_*B*_ is rejected in favor of the alternative *H*_1_ : *µ*_*A*_ ≠ *µ*_*B*_, if *p <* 0.05. We found that, when setting *µ*_*A*_ = *µ*_*B*_ = 3, such that *H*_0_ is true, all approaches considered have a false positive rate (FPR) of around 5% (results not shown^9^) for all levels of trial collinearity (*t*_isi_) and signal-to-noise ratio (*σ*^2^). Therefore, none of the approaches inflates the FPR beyond its nominal level. Furthermore, when setting 5 = *µ*_*A*_ ≠ *µ*_*B*_ = 3, such that *H*_1_ is true, the true positive rate (TPR) of LS-A drastically suffers from a combination of short stimulus intervals and high noise variance (see Figure 4, lower-left panel) whereas ITEM reaches or outperforms LS-S in terms of power (see Figure 4, lowermost row). Therefore, ITEM offers the most powerful test across all scenarios considered.

**Figure 4.**
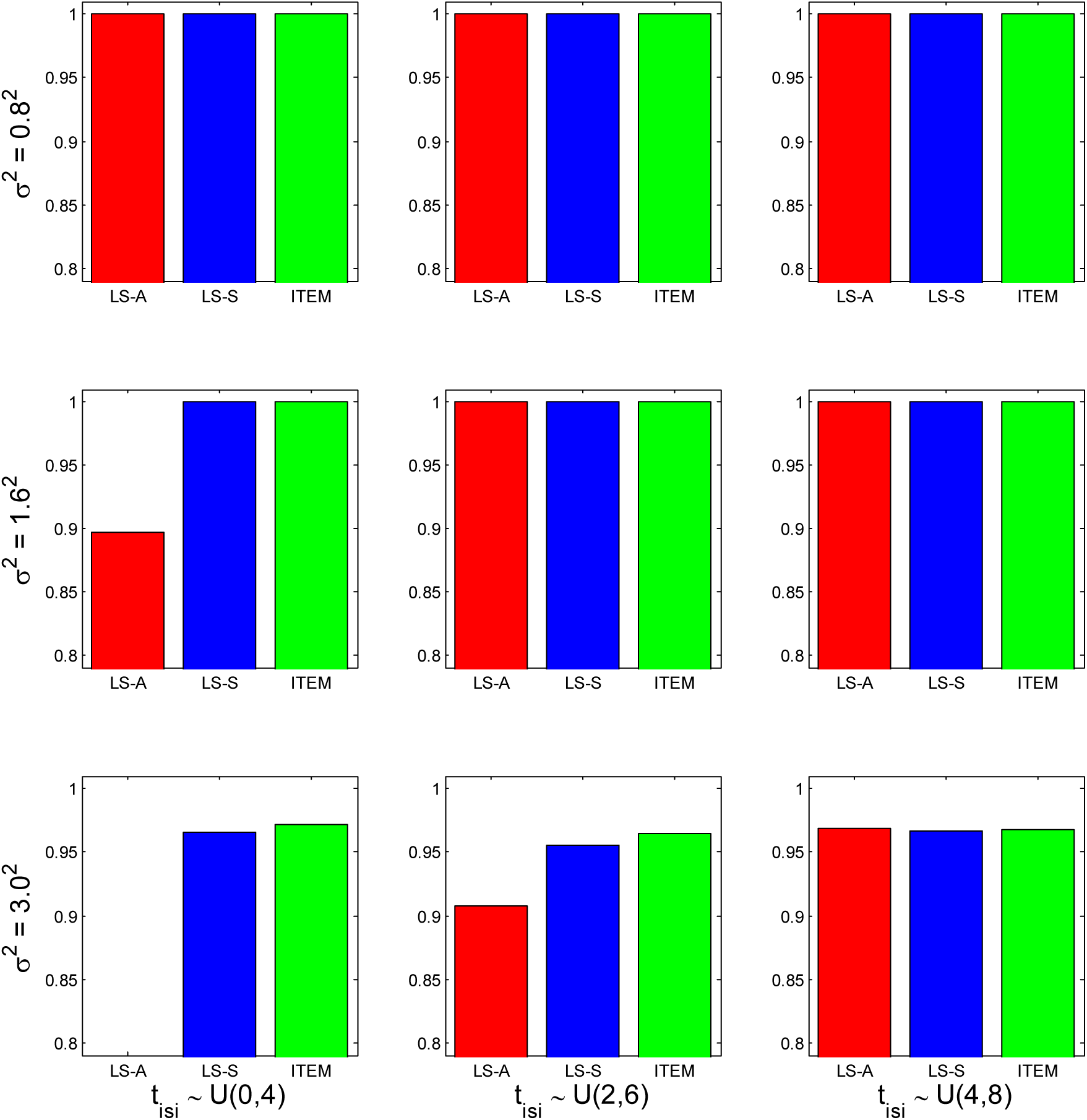
Simulation validation: statistical power. For each combination of inter-stimulusintervals (*t*_isi_) and noise variance (*σ*^2^), the true positive rate (TPR) of a two-sample t-test between trial-wise response amplitudes from two experimental conditions is given for the naïve approach (LS-A, red), the standard approach (LS-S, blue) and the proposed approach (ITEM, green). For long *t*_isi_ and low *σ*^2^, all tests have power of 100%. When the noise variance is high (bottom row) or inter-stimulus-intervals are short (left column), the ITEM approach outperforms or levels with the state-of-the-art approach.

Of course, the two experimental conditions cannot only be statistically tested against each other, but also read out from the generated data. A very simple method for decoding between two conditions is logistic regression. For LS-A and LS-S, condition labels for A and B are coded as 1 and 2 and the corresponding logistic model is estimated. Then, log-odds for the left-out session are predicted from trial-wise response amplitudes and trials are classified into conditions A and B (Mumford et al., 2012). For ITEM, as the presence of correlations in trial-wise parameter estimates makes logistic regression more difficult, the decoding procedure outlined above (see Sections 2.8 and 2.9) was employed for cross-validated classification of trial types. For all approaches, proportion correct (*P*_corr_), i.e. the percentage of trials correctly assigned, was used as the measure of decoding accuracy and decoding accuracy was averaged over the two sessions.

Each procedure leads to one *P*_corr_ per simulation, the distributions of which are visualized as box plots across simulations. We found that, when setting *µ*_*A*_ = *µ*_*B*_ = 3, such that no difference between the conditions exist, all approaches considered have an average decoding accuracy of around 50% (results not shown^9^) for all levels of trial collinearity (*t*_isi_) and signal-to-noise ratio (*σ*^2^). Therefore, there is no evidence for above-chance classification in the absence of a real effect. Furthermore, when setting 5 = *µ*_*A*_ ≠ *µ*_*B*_ = 3, such that there is a real effect, LS-A drastically suffers from a combination of short stimulus intervals and high noise variance (see Figure 5, lower-left panel), whereas median decoding accuracies of ITEM are up to 8.33% higher than those of LS-S (see Figure 5, upper-left panel), but at most 0.83% smaller than them (see Figure 5, lower-left panel). Therefore, the ITEM approach outperforms the original simulation’s best approach in terms of sensitivity, when considered across simulation scenarios.

**Figure 5.**
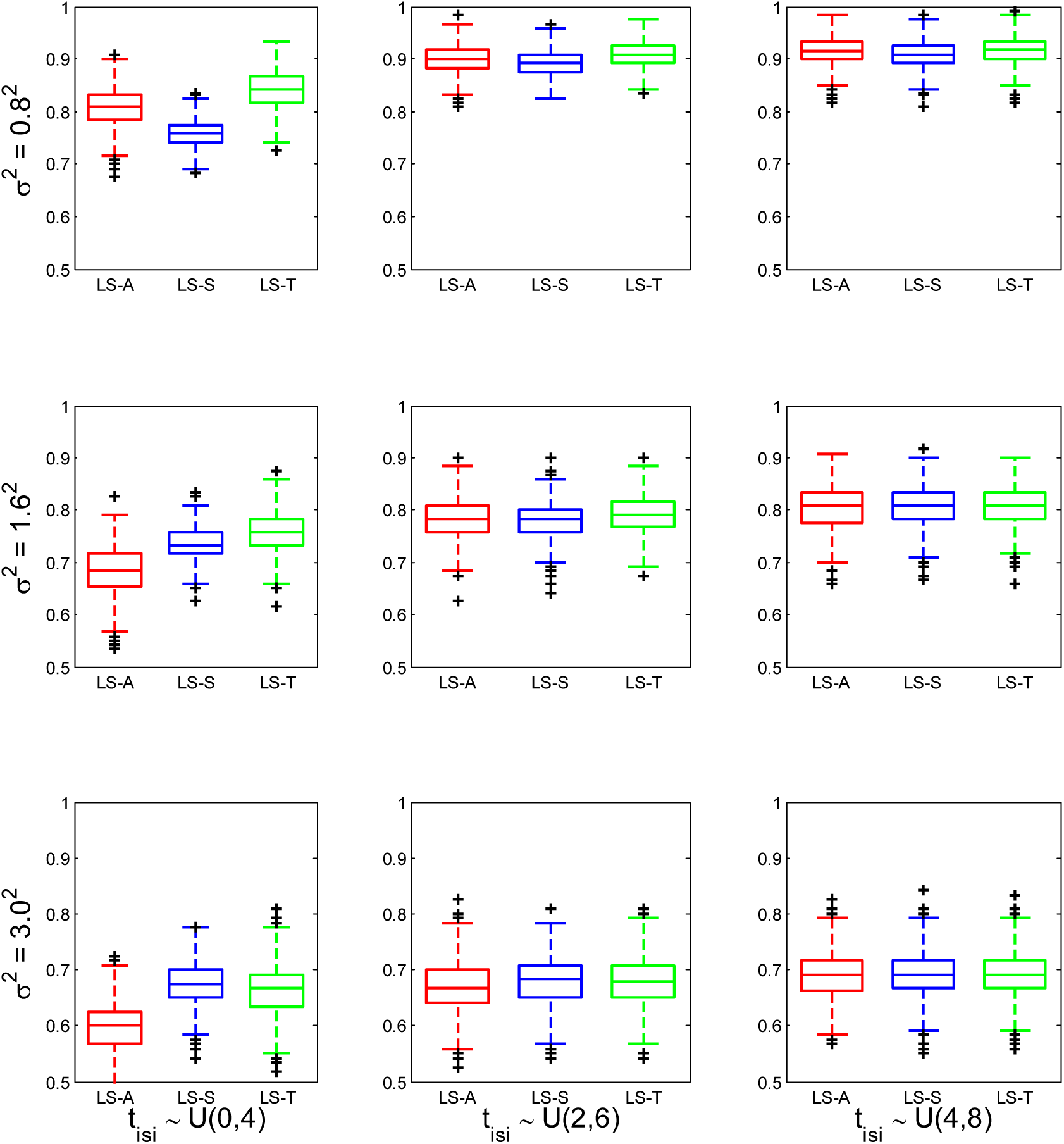
Simulation validation: decoding accuracies. For each combination of interstimulus-intervals (*t*_isi_) and noise variance (*σ*^2^), decoding accuracies for classification between two experimental conditions are given for the naïve approach (LS-A, red), the standard approach (LS-S, blue) and the proposed approach (ITEM, green). For long *t*_isi_ and low *σ*^2^, decoding accuracies of all algorithms are close to 1. When the noise variance is high (bottom row) or inter-stimulus-intervals are short (left column), the ITEM approach outperforms or levels with the state-of-the-art approach. In each boxplot, the central mark is the median; the box edges are the 25th and 75th percentiles; whiskers correspond to the most extreme data points within 1.5× the interquartile range from the box edges; and black crosses represent outliers.

## 4 Application

### 4.1 Experiment

For empirical validation, we re-analyze data from an earlier experiment on visual receptive fields (Heinzle et al., 2011) that was designed to investigate relationships between sensory-visual and cortico-cortical receptive fields. This data set is available in BIDS format from OpenNeuro (https://openneuro.org/datasets/ds002013).

Four right-handed, healthy subjects participated in a retinotopic mapping experiment that was used to define regions of interest (ROI) for visual cortices (V1, V2, V3 and V4, separately for left and right hemisphere) as well as a visual stimulation experiment that is used for ITEM-based visual reconstruction in the present work.

In the main experiment, subjects were viewing a dartboard-shaped flickering checkerboard stimulus (see Figure 6A). The whole display was subdivided into 4 rings and 12 segments, giving rise to 48 sectors (see Figure 6B). Across trials, these sectors changed their local contrast independently and randomly between 4 levels generated using M-sequences (Buračas and Boynton, 2002). These intensity levels were logarithmically spaced between 0.1 and 1 and used for analysis as linearly spaced between 0 and 1 in steps of 1/3. The duration of one trial was 3 s and there was no inter-stimulus-interval (*t*_isi_ = 0 s). In total, 100 trials were presented in each of the 8 sessions. In addition, there was a 15 s rest period at the beginning and at the end of each session.

**Figure 6.**
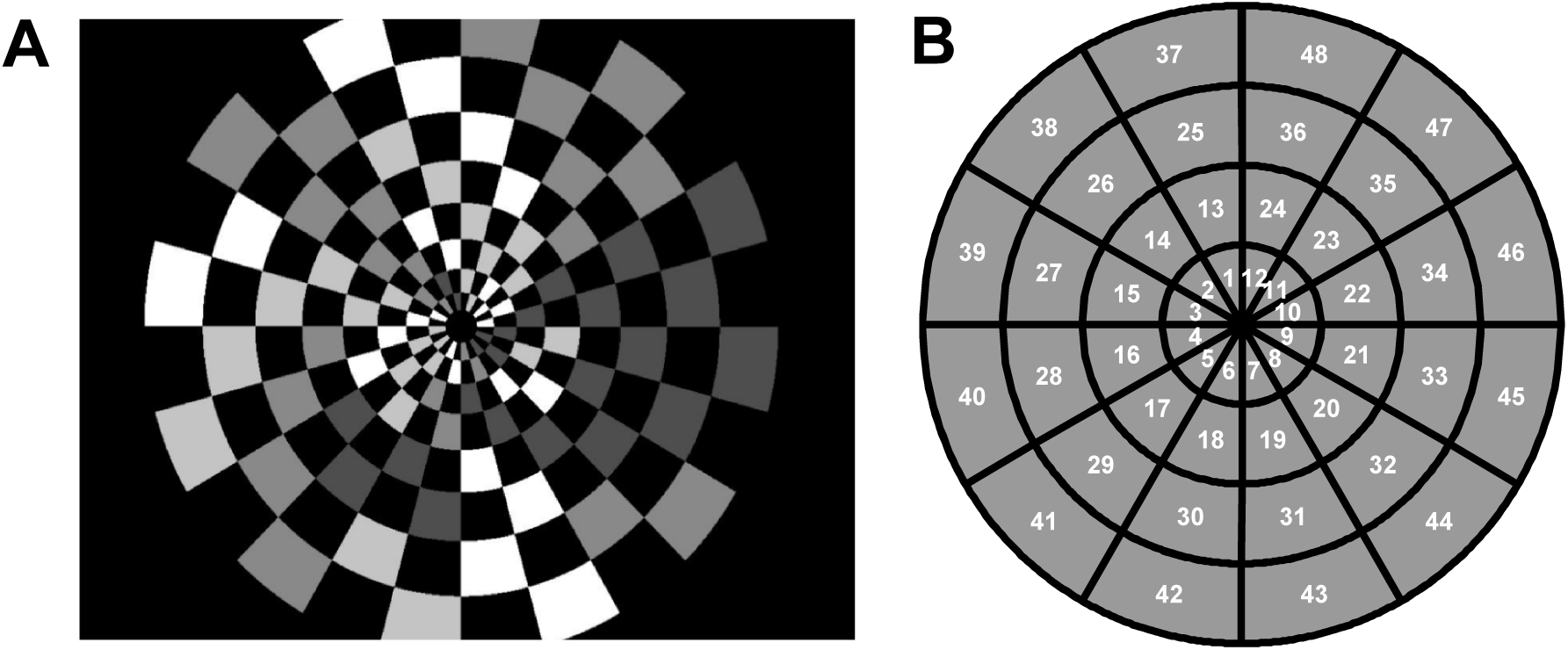
Empirical validation: experimental paradigm. (A) Exemplary stimulus display during a single trial of the receptive field mapping experiment. (B) Schematic view of the 48 sectors which the stimulus display consists of. Each numbered field on the right corresponds to a 2 × 2 checkerboard on the left.

Throughout the experiment, subjects were engaged in a control task to keep their fixation to the center of the visual display. Landolt’s C was presented in the middle of the screen and subjects had to indicate whether it opened to left or to the right side. The open and close times were 800 ms each, with a total stimulus duration of T = 1.6 s in order to avoid divisibility with the acquisition TR = 1.5 s.

Magnetic resonance imaging (MRI) data were collected on a 3-Tesla Siemens Trio with a 12-channel head coil. In each session of the visual stimulation experiment, 220 T2*-weighted, gradient-echo EPIs were acquired at a repetition time TR = 1,500 ms, echo time TE = 30 ms, flip angle *α* = 90° in 25 slices (slice thickness: 2 mm (+ 1 mm gap); matrix size: 64 × 64) resulting in a voxel size of 3 × 3 × 3 mm. During the separate retinotopic mapping experiment, 160 T2*-weighted volumes were acquired with 33 slices, TR = 2,000 ms and all other parameters as above.

### 4.2 Analysis

In pre-processing, fMRI data were converted from 3D into 4D NIfTIs, transformed into the BIDS format (Gorgolewski et al., 2016), reoriented to the axis from commissura anterior (AC) to commissura posterior (PC), acquisition-time-corrected (slice timing) and head-motion-corrected (spatial realignment) using SPM12. The retinotopic mapping was based on a standard traveling wave method (Wandell et al., 2007) and analyzed to yield flattened angular and eccentricity maps (Heinzle et al., 2011).

In statistical analysis, an ROI-based ITEM analysis was performed via these steps:

1. *standard GLM specification*: A standard design matrix *X* was specified with the following regressors: (i) 1 onset regressor describing visual stimulation throughout the experiment; (ii) 48 parametric modulators describing intensity levels in the 48 sectors of the visual stimulus; (iii) 2 onset regressors describing the control fixation task during the experiment; (iv) 6 motion regressors describing head movements; (v) 5 filter regressors describing periodic drifts; and (vi) 1 constant regressor describing the implicit baseline. An exemplary design matrix for one session from one subject is given on the left of Figure 1B.
2. *standard GLM estimation*: Using this model, the temporal covariance matrix *V* was estimated using SPM’s AR(1) model. An exemplary covariance matrix for the same session and subject is given in the middle of Figure 2B.
3. *trial-wise GLM specification*: Using SPM-compatible MATLAB code (see Section 6), the trial-wise design matrix *X*_*t*_ (see Figure 1B) and the transformation matrix *T* were generated based on design information assembled during SPM model specification.
4. *trial-wise GLM estimation*: Trial-wise parameter estimates 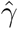 and the trial-by-trial covariance matrix *U* (see Figure 2B) were calculated using equations (4) and (7).
5. *model comparison* & *voxel selection*: As our goal was not searchlight decoding, but visual reconstruction from V1, we performed an intermediate step^10^ of cross-validated Bayesian model selection (cvBMS; Soch et al., 2016) using routines from the MACS toolbox (Soch and Allefeld, 2018) to identify voxels processing visual information:
  a. To this end, 48 single-sector models, each describing intensity levels in one of the 48 sectors, and 1 null model, describing no individual sector, were specified as design matrices *T* predicting trial responses 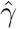 (see eq. 8). Then, models were assessed using the cross-validated log model evidence (cvLME) and the family of single-sector models was compared against the null model.
  b. In each hemisphere, the 48 V1 voxels with the highest evidence in favor of the family of single-sector models were identified, resulting in a combined ROI containing 96 voxels used for reconstruction in the left-out session.
6. *reconstruction*: Finally, session-wise transformation matrices *T*, uncorrelation matrices *U* and response amplitudes within the selected voxels 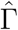 were used for cross-validated reconstruction of intensity levels in the 48 sectors using equations (14) and (15). For all reconstructions, decoding accuracy was quantified via correlation coefficients and then averaged over sessions and subjects, but not sectors.

The complete empirical data analysis can be reproduced using MATLAB code available from GitHub (https://github.com/JoramSoch/ITEM-paper).

### 4.3 Results

The present experiment constitutes an ideal proof of concept for the ITEM framework due to (i) the extremely rapid event-related design without any inter-stimulus-intervals and (ii) the massive parametric design information to be decoded.

Using an ITEM-based analysis, visual contrast in almost all parts of the visual field could be reliably decoded from fMRI signals in left and right V1 (see Figure 7C). Reconstruction performance is better for sectors which are far from the center (e.g. 45 vs. 9) and for sectors which are close to the horizontal axis (e.g. 45 vs. 48) of the visual field, providing evidence for the so-called “oblique effect” in visual cortex (Li et al., 2003).

**Figure 7.**
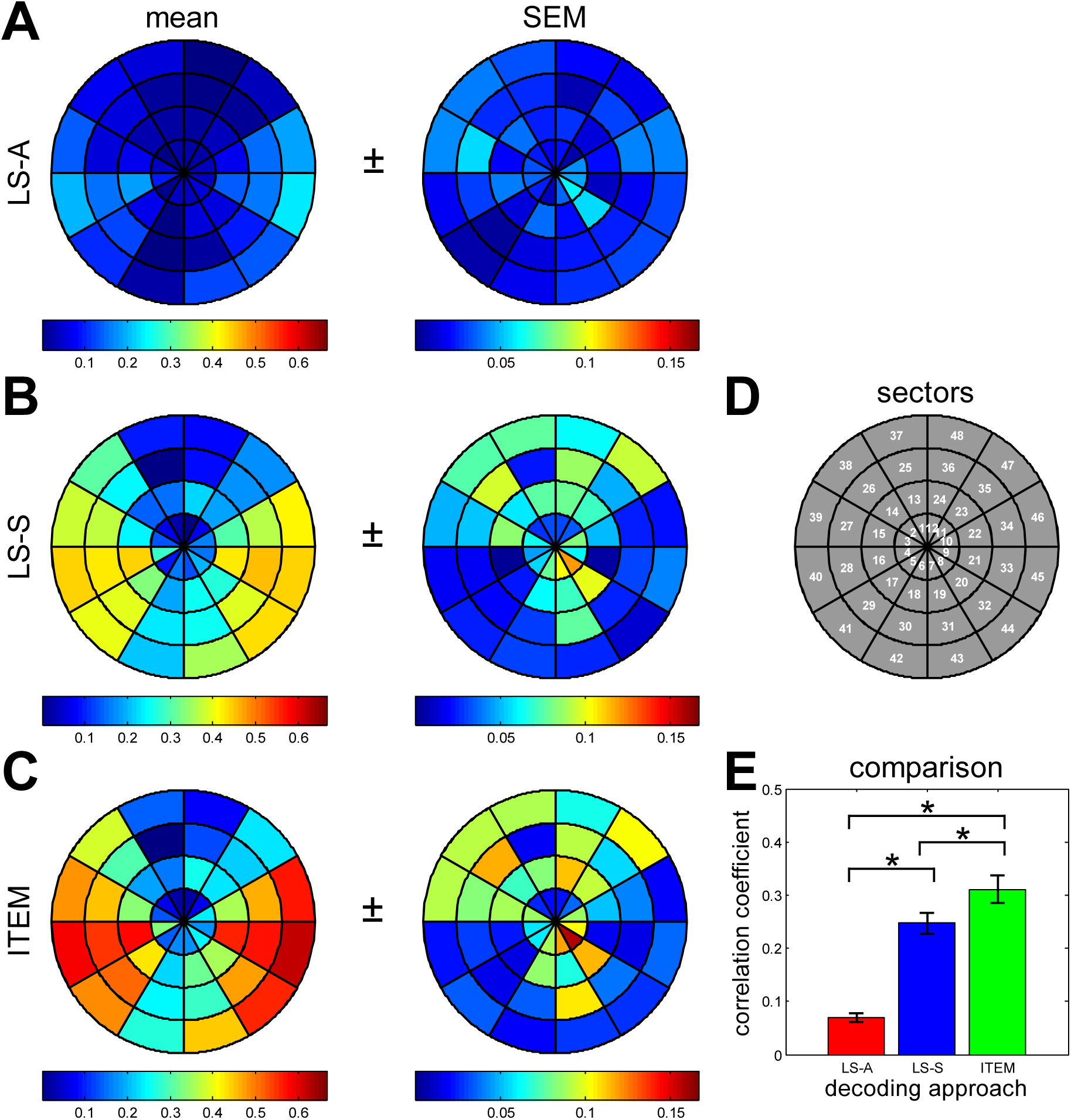
Empirical validation: decoding accuracies. For each sector, cross-validated decoding accuracies are given as correlation coefficients between presented contrast and reconstructed contrast, when using (A) the naïve approach ignoring trial-by-trial correlations (LS-A), (B) the standard approach using separate design matrices (LS-S) and (C) the proposed approach, i.e. inverse transformed encoding modelling (ITEM). Each row shows mean correlations, averaged over 8 sessions and 4 subjects (left column), as well as standard errors over subjects (middle column). For comparison purposes, all panels in one column use the same color axis. (D) For orientation purposes, the sector indices are recapitulated. (E) The three approaches are summarized using average mean correlations and standard errors over sectors (lower-right panel). ITEM-based reconstruction (mean *r*: 0.311) strongly outperforms LS-A (mean *r*: 0.070) and mildly outperforms LS-S (mean *r*: 0.247). All pairwise comparisons are significant at *p <* 0.001.

To compare the ITEM approach against LS-A and LS-S, the same reconstruction was applied without incorporating the temporal covariance matrix *U* (LS-A) and to trial-wise response amplitudes estimated via separate design matrices *X*_*i*_ (LS-S).^11^ As expected, LS-A provided the lowest correlation coefficients when compared to LS-S or ITEM (see Figure 7A), probably suffering from high trial-by-trial correlations due to the short interstimulus-intervals. LS-S improves significantly over LS-A in terms of decoding accuracies (see Figure 7B), but is still outperformed by ITEM (see Figure 7E).

From the correlation coefficients across all subjects, sessions and sectors obtained with ITEM-based reconstruction, the highest correlation, the lowest absolute correlation and the median correlation were selected as examples for a particularly good, a particularly bad and a medium-quality reconstruction (see Figure 8). Time courses of presented contrast and reconstructed contrast can be plotted with each other, showing considerable covariation for the best reconstruction (see Figure 8C).

**Figure 8.**
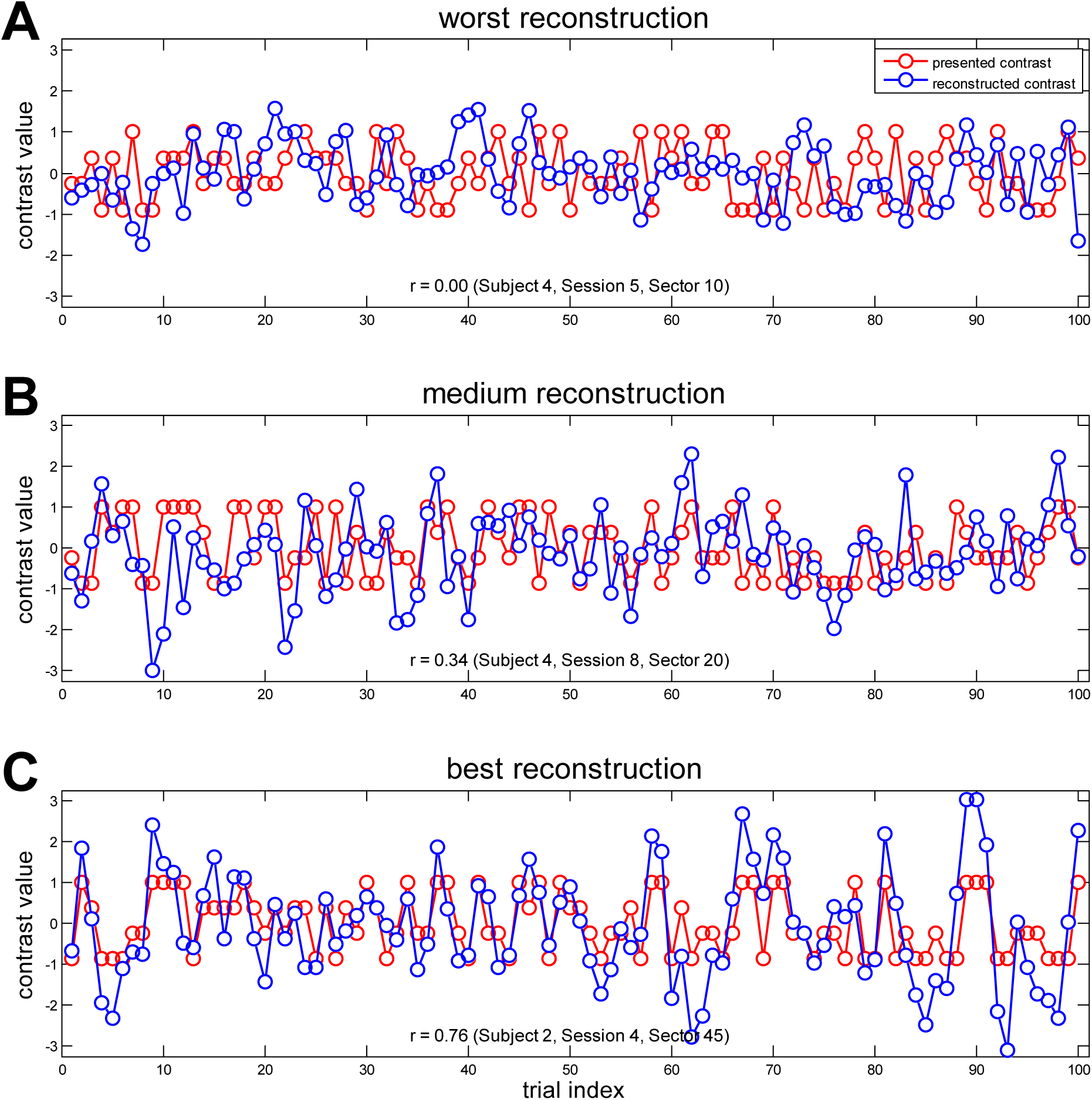
Empirical validation: exemplary reconstructions. From all ITEM-based decoding analyses (see Figure 7C), (A) the worst reconstruction, corresponding to the lowest absolute correlation coefficient, (B) a medium reconstruction, corresponding to the median among all correlations, and (C) the best reconstruction, corresponding to the highest correlation coefficient, were selected for display. Each panel shows the actually presented contrast (red), normalized to a range from −1 to +1, and the reconstructed contrast (blue) across trials.

## 5 Discussion

We have introduced *inverse transformed encoding modelling* (ITEM), an integrated framework for trial-wise linear decoding of experimental manipulations from fMRI data that accounts for trial-by-trial correlations and thus avoids suboptimal decoding accuracies resulting from inaccurate parameter estimates. ITEM allows for classification of discrete experimental conditions as well as reconstruction of continuous modulator variables. In a simulation study on binary classification, ITEM reached or outperformed LS-S, the previously best known approach (Mumford et al., 2012), for most noise variances and inter-stimulus-intervals tested (see Figure 5). In an empirical application to visual reconstruction, ITEM was used to successfully decode visual stimulation from multivariate signals in left and right V1 (see Figure 7).

The problem of *correlated trial-by-trial parameter estimates* has already been discussed several times in the fMRI/MVPA literature (Mumford et al., 2012; Turner et al., 2012; Mumford et al., 2014; Weeda, 2018). Previous contributions have pointed out that a naïve approach ignoring trial-by-trial correlations (i.e. LS-A) leads to suboptimal parameter estimates and found that a revised method estimating each trial’s activation using a separate design matrix (i.e. LS-S) better controls for trial-to-trial covariance. This procedure is based on the idea that including a regressor modelling all other trials in addition to the regressor modelling the trial of interest will largely reduce collinearity between trials (Mumford et al., 2012).

Here, we extend this previous work by providing a principled approach, based on the actual *distribution of the trial-wise parameter estimates* (i.e. ITEM), as implied by the trial-wise design matrix (see Section 2.6 and Equation 7; see also Mumford et al., 2014, p. 132). Unlike LS-S, ITEM is not simply an alternative way of calculating trial-wise parameter estimates, but an integrated decoding approach accounting for trial-by-trial correlations (see Section 2.8).

However, ITEMs are not only applicable to rapid event-related designs, but generally useful when *fMRI-based trial-wise linear decoding* is the goal. Other than a decoding algorithm, e.g. a linear SVM applied to trial-wise response amplitudes, ITEM controls for correlation of trial-specific activations with any covariate present in the experimental design (see Figure 2B and Equation 6). A disadvantage compared to other approaches (e.g. Weeda, 2018) is that ITEM requires an assumption about the shape of the hemodynamic response in the form of an HRF (e.g. the canonical HRF). Future research may investigate whether the (inverse) transformation encoding model can be combined with a trial-wise GLM using a finite impulse response (FIR) basis set in order to allow for more flexibility with respect to the HRF shape.

The ITEM approach is very similar to the technique of *inverted encoding models* (IEM; Sprague et al., 2015) that is used frequently for reconstruction of sensory information (Brouwer and Heeger, 2009; Saproo and Serences, 2014). In Appendix D, we outline two key differences between ITEMs and IEMs, namely (i) the reversed order of model estimation and model inversion in the reconstruction process and (ii) the fact that IEM in its most frequent implementation does not account for possible covariance between trials by not considering the *U* matrix.

ITEM has been made available as an *SPM plug-in* on GitHub (see Section 6). While the present work only used decoding from regions of interest (ROI), an implementation for decoding from searchlights (SL) is also available. We hope that ITEM-based decoding will increase the sensitivity of MVPA for rapid event-related fMRI designs.

## 6 Implementation

An SPM12-compatible MATLAB implementation of the ITEM approach (https://github.com/JoramSoch/ITEM) as well as all code underlying the analyses in this paper (https://github.com/JoramSoch/ITEM-paper) are available from GitHub.

The data set used for empirical validation in Section 4 has been BIDS-formatted and uploaded to OpenNeuro (https://openneuro.org/datasets/ds002013). Further instructions on data processing can be found in the readme file of the accompanying repository (https://github.com/JoramSoch/ITEM-paper/blob/master/README.md).

## 7 Acknowledgements

This work was supported by the Bernstein Computational Neuroscience Program of the German Federal Ministry of Education and Research (BMBF grant 01GQ1001C), the Collaborative Research Center “Volition and Cognitive Control: Mechanisms, Modulations, Dysfunctions” (SFB 940/1) and the German Research Foundation (DFG grants EXC 257 and KFO 247).

The authors would like to thank Jakob Heinzle (TNU/ETH Zürich) for acquiring the fMRI data set used for empirical validation and for permission to make this data set publicly available.

The authors have no conflict of interest, financial or otherwise, to declare.

## 8 Appendix

## A Derivation of the transformed encoding model

The following derivation of the transformed encoding model (TEM) makes use of the *linear transformation theorem* for the *multivariate normal distribution*:

### Theorem 0

Let *x* follow a multivariate normal distribution:

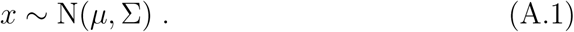

Then, any linear transformation of *x* is also multivariate normally distributed:

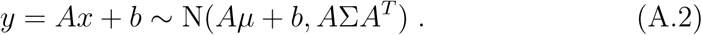

**Proof:** This is standard textbook knowledge (see e.g. Koch, 2007, eq. 2.202).

To recapitulate (see Sections 2.2–2.3), the standard general linear model (GLM) and the trial-wise GLM for first-level fMRI data analysis are given by

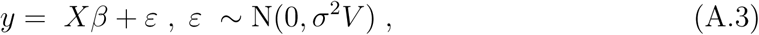

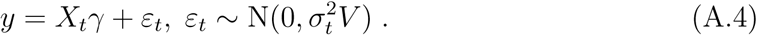

These two models are linked to each other by the transformation matrix (see Section 2.4):

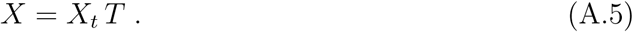

Parameter estimates for the trial-wise GLM are given by weighted least squares:

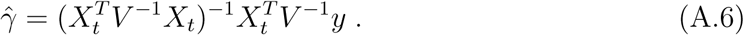

The distribution of these parameter estimates is specified by the *1st TEM theorem*:

### Theorem 1

The trial-wise parameter estimates 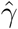 are distributed as

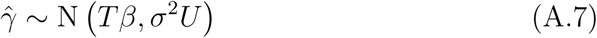

when (A.3) is the ground truth, where the covariance matrix *U* is given by

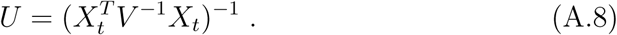

**Proof:** Combining (A.3) with (A.2), we have:

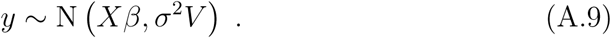

Note that the 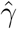 given in (A.6) is a linear transformation of the *y* given by (A.9). Thus, we can again apply (A.2) which gives:

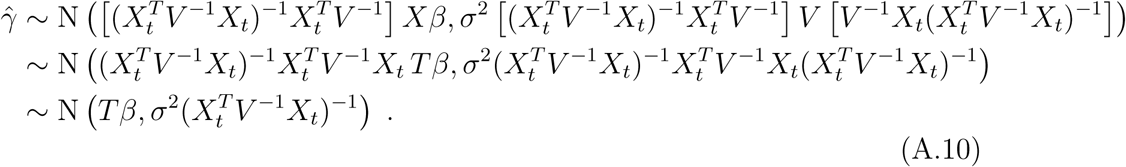

This distribution can also be written as the following equation which is referred to as the *transformed encoding model* (TEM):

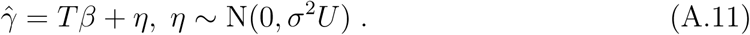

The *U* matrix is an important component of the transformed encoding model framework, because it accounts for the covariance induced into the trial-wise parameter estimates by overlapping HRFs in rapid event-related designs. Being derived from the trial-wise design matrix (A.4), it simply adjusts for the correlations caused by this estimation method in the trial-wise encoding model (A.11). Parameter estimates for this model are again given by weighted least squares:

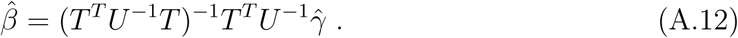

An equivalence of these parameter estimates is stated by the *2nd TEM theorem*:

### Theorem 2

The parameter estimates of the TEM

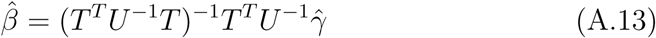

are equivalent to the parameter estimates of the standard GLM

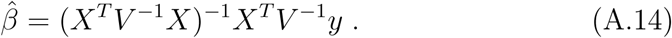

**Proof:** To see this, apply the inverse covariance matrix from (A.8), the transformation matrix definition in (A.5) and the trial-wise parameter estimates given by (A.6) to the condition-wise parameter estimates given by (A.12):

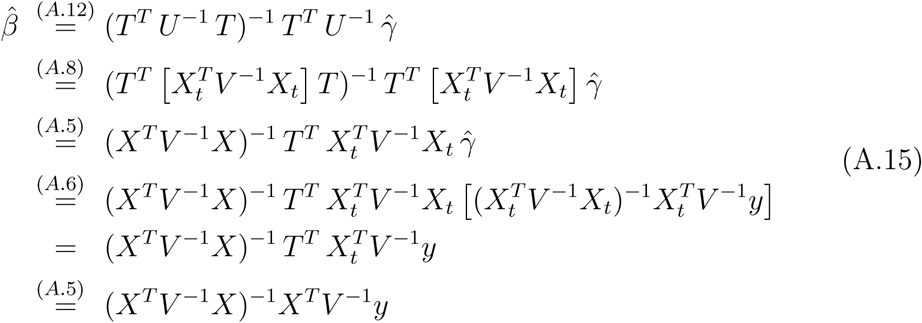

This demonstrates that parameter estimates from the trial-wise model (A.11) are equivalent to parameter estimates from the scan-wise model (A.3) when the transformation matrix *T* is chosen in a way that maps from the trial-wise design matrix *X*_*t*_ to the standard design matrix *X* (see Figure 1). This is achieved by virtue of the trial-by-trial covariance matrix *U* that is derived from *X*_*t*_ to accommodate the correlation introduced into trial-wise parameter estimates 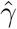 by using HRFs overlapping in time (see Figure 2).

## B Extension with trial-wise response variability

Note that, when proving Theorem 1, *y* was assumed to be generated by the standard GLM (A.3) assuming equal trial-wise response amplitudes within experimental conditions. In contrast, if *y* is assumed to be generated by the trial-wise GLM (A.4) allowing for variation of trial-wise response amplitudes within experimental conditions, the distribution of 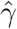 becomes more complicated, as asserted by the *3rd TEM theorem*:

### Theorem 3

When the true model is the trial-wise GLM

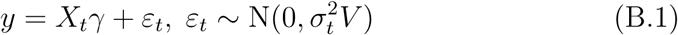

and the trial-wise response amplitudes *γ* follow the equation

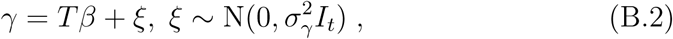

then the trial-wise parameter estimates 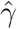 from (A.6) are distributed as

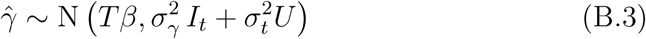

where 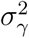 is the trial-to-trial variance and *U* is given by (A.8).

**Proof:** When the trial-wise GLM is true, this means that response amplitudes differ across trials which is equivalent to the assumption that the trial-to-trial variance 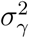 is not zero. Here, we make the assumption that the true trial-wise response amplitudes are drawn as i.i.d. samples with an expectation that is a function of the trial matrix *T* and condition-wise activations *β*, as indicated by (B.2). Together with (B.1) and (A.5), this implies:

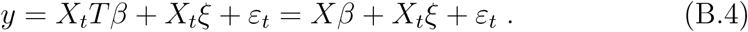

Applying (A.2) to (B.4) and summing the covariances of the independent normal variates *X*_*t*_*ξ* and *ε*_*t*_, it follows that (cf. Mumford et al., 2014, eq. 4):

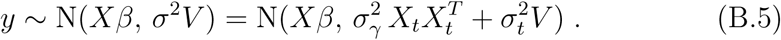

The 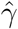 given in (A.6) is a linear transformation of the *y* given by (B.5). Thus, we can again apply (A.2) which gives (cf. Mumford et al., 2014, eq. 6):

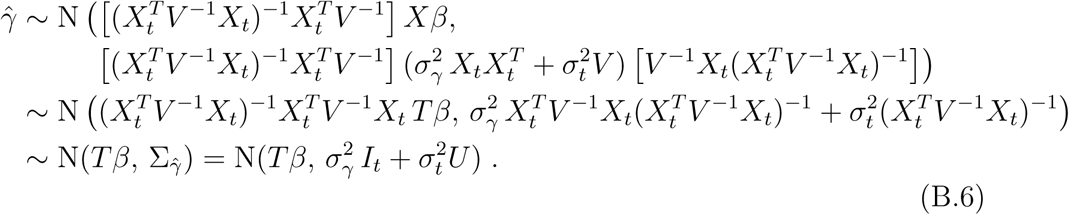

Formally, with moving from (A.4) to (B.4), the first-level GLM changes from a fixed-effects model into a random-effects model, because *γ* becomes a random variable by (B.2). This also allows for a new interpretation of (A.3), since its covariance *σ*^2^*V* is now replaced by two components, as given in (B.5).

As becomes apparent from (B.6), the covariance of the trial-wise parameter estimates also consists of two components, one coming from the original trial-to-trial variability assumed by (B.2) (the “natural” covariance) and the other due to the trial-wise design matrix *X*_*t*_ via *U* (the “induced” covariance).

If the trial-to-trial variance is assumed to be zero, i.e. 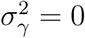, the distribution becomes

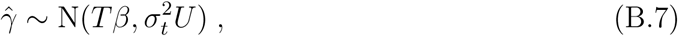

such that the model can be estimated via *weighted least squares* (WLS):

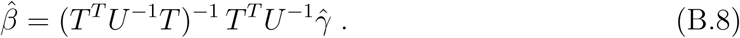

On the other hand, if inter-stimulus-intervals are sufficiently long^12^, such that trial-wise HRF regressors are not correlated, then *U* = *u*_*t*_ *I*_*t*_ and the distribution becomes

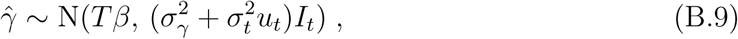

such that the model can be easily estimated via *ordinary least squares* (OLS):

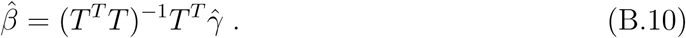

However, in the presence of covariates (see Figure 2B) or in rapid event-related designs (see Figure 2A), this is practically never fullfilled so that *U* ≠ *u*_*t*_ *I*_*t*_ and a *variance components model* (VCM; Searle et al., 1992) with known covariance components (*I*_*t*_, *U*) and unknown variance factors 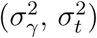 has to be estimated.

Let 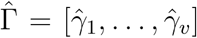 be a *t* × *v* matrix of horizontally concatenated trial responses over voxels with number of voxels *v*. Then, the VCM in (B.3) can be inverted using SPM’s restricted maximum likelihood (ReML) algorithm (Friston et al., 2002a) which has to be implemented via the SPM command

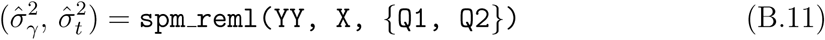

where 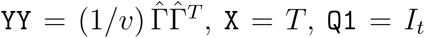, `X` = *T*, `Q1` = *I*_*t*_ and `Q2` = *U*. The function output can then be used to calculate ReML estimates as

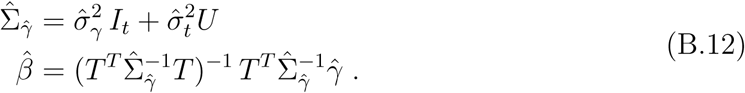

In principle, ReML estimation can be performed when *v* = 1, but estimates of variance components become more accurate with increasing number of signals, i.e. voxels. In our simulation study, we have used (B.11) with *v* = 1,000 signals to improve ITEM-based classification (see Appendix E). In our empirical validation, we have found no improvement of ITEM-based reconstruction by ReML estimation which is why this extension is currently not implemented in the released code (see Section 6).

## C Conversion from forward to backward model

Extending the univariate transformation encoding model (TEM) from (A.11) to several voxels, one arrives at the multivariate TEM (see Section 2.7) which also forms the basis for motivating the inverse TEM (see Section 2.8). This way of proceeding from the MTEM to the ITEM is the subject of the *4th TEM theorem*:

### Theorem 4

Consider the multivariate transformed encoding model

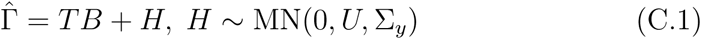

and let *B W* = *I*_*p*_. This implies the inverse transformed encoding model

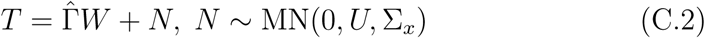

with the covariance matrix Σ_*x*_ = *W* ^*T*^ Σ_*y*_*W*.

**Proof:** If there is a *v* × *p* matrix *W* such that *B W* = *I*_*p*_, this extraction filter *W* is the right-inverse of the activation pattern *B* (cf. Haufe et al., 2014, eq. A.1). Such a matrix exists, if the rows of *B* are linearly independent or, in other words, if all regressors in *T* have mutually dissociable activation patterns. Then, right-multiplying the multivariate forward model with *W* yields

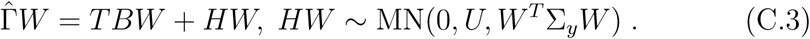

Applying *B W* = *I*_*p*_ and rearranging, we have

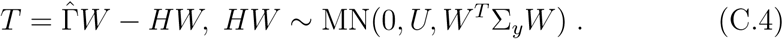

Substituting *N* = *HW* and Σ_*x*_ = *W* ^*T*^ Σ_*y*_*W*, we get

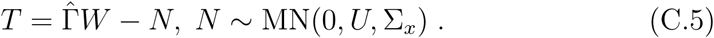

Because *N* is mean-free and a zero-mean (matrix-)normal distribution is symmetric around zero, -*N* has the same distribution as +*N*, such that

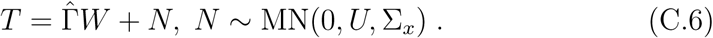

Note that in step (C.3), because we are multiplying with *W* from the right and not from the left, Theorem 0 (see eq. A.2) does not affect the temporal covariance *U*, but has to be applied to the spatial covariance Σ_*y*_. In fact, because the dependent variable in (C.6) is *T*, this is not a spatial covariance anymore, but rather a design covariance now, as it pertains to correlations between trial-wise experimental design variables.

This makes the new model somewhat problematic, because other than *B* in (C.1), the parameter matrix *W* enters in (C.2) in two ways: as the mapping from 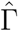 to *T* and in the distribution of the noise matrix *N*. However, if the estimation of *W* can be understood as separate column-by-column operations on *T*, this problem can be ignored as Σ_*x*_ only applies to relations between columns of *T*. In fact, the estimates used for reconstruction in (14) are in some sense optimal, as stated by the *5th TEM theorem*:

### Theorem 5

Given the inverse transformed encoding model

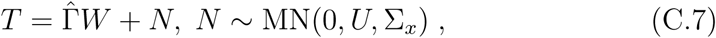

the weighted least squares solution for the weight matrix

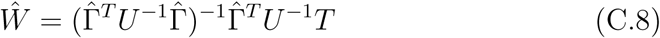

is the best linear unbiased estimator (BLUE) of *W*.

**Proof:** The best linear unbiased estimator 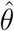 of a certain quantity *θ* estimated from measured data *y* is 1) an estimator resulting from a linear operation *f* (*y*), 2) whose expected value is equal to *θ* and 3) which has, among those satisfying 1) and 2), the minimum variance.

1. First, Ŵ is a *linear* estimator, because it is of the form 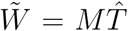 where *M* is an arbitrary *v* × *t* matrix.
2. Second, Ŵ is an *unbiased* estimator, if ⟨*Ŵ*⟩ = *W*. By applying (A.2) to (C.7), the distribution of 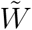 is

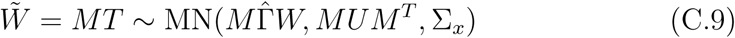

which requires that 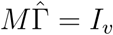. This is fulfilled by any matrix 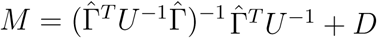 where *D* is a *v* × *t* matrix which satisfies 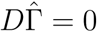.
3. Third, the *best linear unbiased* estimator is the one with minimum variance, i.e. the one that minimizes the expected Frobenius norm

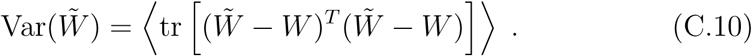 Using the distribution of 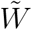 from (C.9)

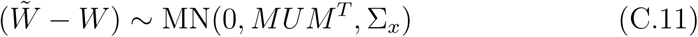

and the Wishart distribution property

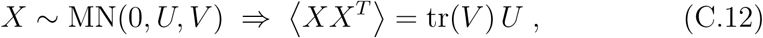

this variance can be evaluated as a function of *M* :

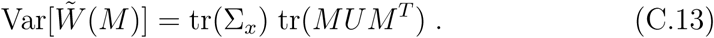 As a function of *D* and using 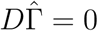, it becomes:

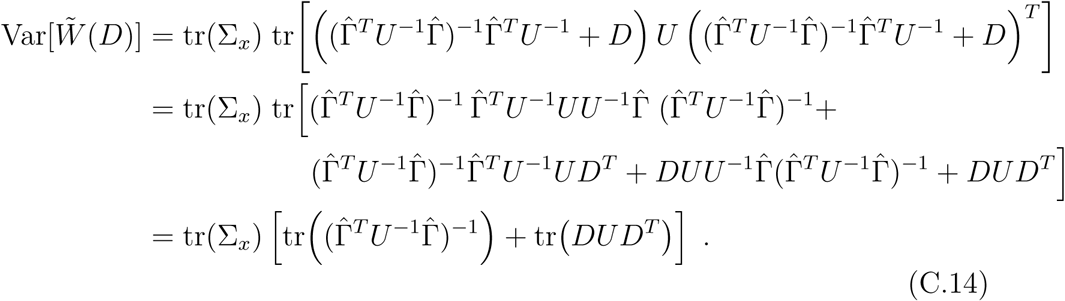

Since *DUD*^*T*^ is a positive-semidefinite matrix, all its eigenvalues are non-negative. Because the trace of a square matrix is the sum of its eigenvalues, the mimimum variance is achieved by *D* = 0, thus producing Ŵ as in (C.8).

## D Relation of ITEMs to inverted encoding models

Let 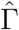 be the *t* × *v* matrix of voxel-and-trial-wise response amplitudes, *T* be the *t* × *p* matrix of trial-wise experimental manipulations and *B* be the *p* × *v* matrix of voxel-wise activation patterns. The following derivation requires that *t* > *v* > *p*, i.e. there are more trials than voxels and there are more voxels than conditions to classify or variables to reconstruct.

Consider the multivariate transformed encoding model (MTEM) given by

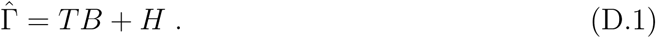

This model can be inverted with respect to the parameter matrix

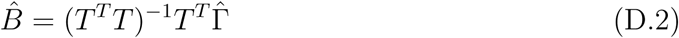

or, when treating *B* as a constant, with respect to the design matrix

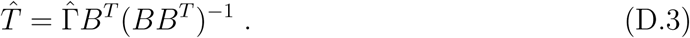

In an inverted encoding model (IEM) analysis, one applies step (D.2) to the training data and then applies step (D.3) to the test data which yields the following reconstruction (e.g. Brouwer and Heeger, 2009, eq. 3; Saproo and Serences, 2014, eq. 3):

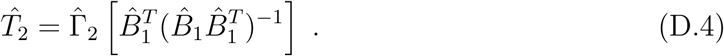

Now consider the inverse transformed encoding model (ITEM) given by

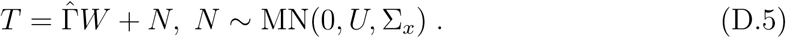

This model can be estimated using weighted least squares

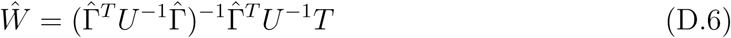

and the predicted, estimated or fitted signals are given by

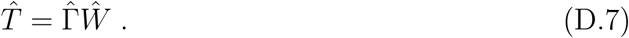

In an ITEM-based analysis, one applies step (D.6) to the training data and then applies step (D.7) to the test data which yields the following reconstruction (see eq. 15):

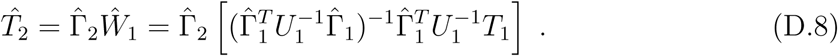

When comparing (D.4) with (D.8), one can see that both of them right-multiply an operator matrix to the test data 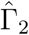, but these matrices are quite different from each other: One difference is that the IEM operator results from inversion of an estimated model whereas the ITEM operator results from estimation of an inverse model. Another and more important difference is that the IEM approach ignores covariance between trials (and covariance of trials with other covariates) indicated by the matrix *U* in (D.6), simply because the parameter matrix is estimated via ordinary least squares in (D.2).

## E Simulation adapted from Mumford et al. (2012)

The generative model underlying Mumford et al.’s simulations can be described as follows. First, in each session from each simulation run, *t* = 60 trials are evenly distributed into 2 experimental conditions or trial types (tt)

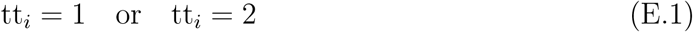

where tt_*i*_ = 1 and tt_*i*_ = 2 for 30 trials each.

Second, trial-wise activations are independently sampled from a normal distribution

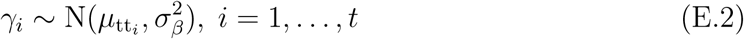

where *µ*_1_ = 5, *µ*_2_ = 3 and 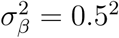.

Third, inter-stimulus-intervals are independently sampled from a uniform distribution

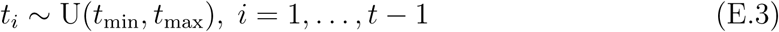

where *t*_min_ ∈ {0, 2, 4} and *t*_max_ = *t*_min_ + 4.

Based on the sampled inter-stimulus-intervals (ISIs), the canonical hemodynamic response function (cHRF) as well as stimulus duration *t*_dur_ and repetition time TR, a trial-wise design matrix *X*_*t*_ is generated which instantiates the sampled ISIs (see Figure 1A as an example for *t*_isi_ ~U(0, 4)). Moreover, a temporal correlation matrix *V* embodying an AR(1) process is generated with *ρ* = 0.12 as

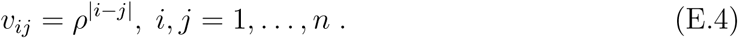

Finally, fMRI signal noise is sampled with standard deviation *σ* ∈ {0.8, 1.6, 3} as

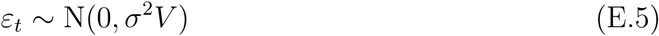

and the simulated data are generated according to the trial-wise GLM as

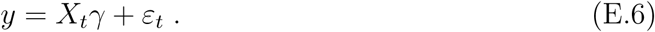

The combination of the 3 different options for *t*_min_/*t*_max_ and the 3 different options for *σ*^2^ leads to 9 different simulation scenarios (see Figures 4 and 5). In each scenario, *N* = 1,000 simulations with *S* = 2 sessions per simulation were performed.

After data generation, trial-wise activations are estimated and trial types are decoded. In the “least squares, all” (LS-A) approach, 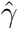 was obtained via equation (4) and a logistic regression was trained on the estimates from one session to predict trial types in the other session and vice versa (Mumford et al., 2012, App. A). Decoding accuracy was quantified as the proportion of trials correctly assigned to trial types 1 and 2 based on calculated log-odds in the test session. In the “least squares, separate” (LS-S) approach, 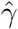 was obtained using a separate design matrix for each trial, including one regressor for this trial and one regressor for all other trials (Mumford et al., 2012, Fig. 1), and the same logistic regression was applied. For inverse transformed encoding modelling (ITEM), 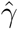 was obtained via equation (4) and trial types were decoded via equations (7), (14) and (15), with decoding accuracy being assessed via equation (17).

The present simulation differs from Mumford et al.’s in the following respects:

- In the original simulation, the simulated data were high-pass filtered although no temporal drifts were in the ground truth. In the present simulation, data were not filtered.
- In the original simulation, the simulated data were not whitened although mild autocorrelation was induced into the ground truth (see Equation E.4). In the present simulation, this was handled by using the *V* matrix in equations (4) and (7).
- These two differences may have contributed to the fact that decoding accuracies reported in the present simulation (see Figure 5) were generally higher than in the original simulation (Mumford et al., 2012, Fig. 3).
- The stimulus duration *t*_dur_ and repetition time TR from the original simulation were not reported in the respective paper and could not be recalled by the corresponding author, so *t*_dur_ = 2 s and TR = 2 s were used.
- The simulation scenarios characterized by *t*_min_ = 6 and *t*_max_ = 10, i.e. with ISIs distributed as *t*_isi_ ~ U(6, 10), were omitted from the simulation, because no changes relative to *t*_isi_ ~U(4, 8) could be observed.
- In the present simulation, the design matrix was identical for all simulation runs belonging to one simulation scenario (as opposed to tt and *t*_isi_ being resampled in every simulation run). This was necessary for applying the ReML approach within the ITEM framework (see Appendix B, esp. Equation B.11) and is consistent with typical fMRI data analysis where the same design matrix (and covariance matrix) is used to analyze all voxels (which correspond to simulation runs).
- This means that the present simulation provides no statistics across randomized designs within a scenario. Nonetheless, repeating the simulation with different random designs gives qualitatively equivalent results.
- In the present simulation, *N* = 1,000 (instead of *N* = 500) simulation runs were used for higher precision in estimating statistical power and decoding accuracies.
- In the present simulation, *S* = 2 (instead of *S* = 3) sessions per simulation were used, because there was no need for the double cross-validation procedure for hyper-parameter tuning required by some other estimation methods considered in the original simulation (Mumford et al., 2012, Fig. 1).

---

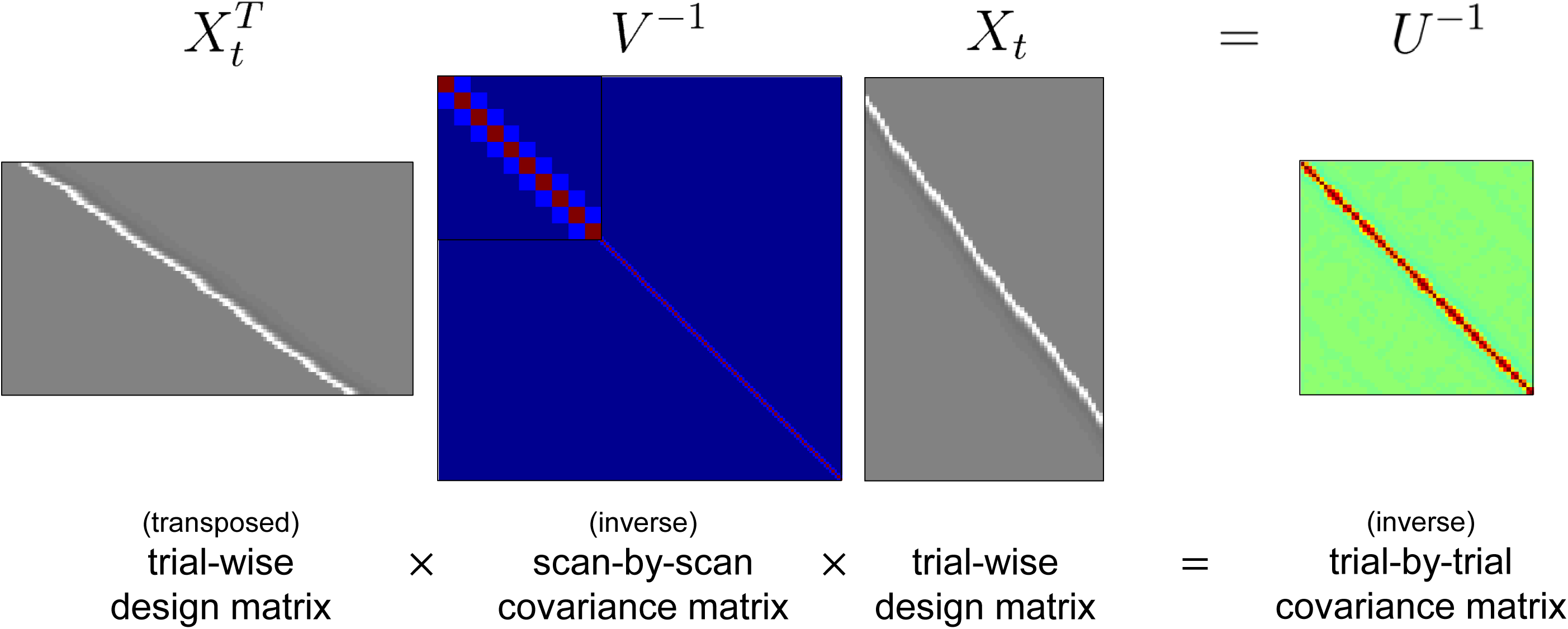

## Footnotes

1 For example, computation time for whole-brain trial-wise parameter estimates in a first-level GLM for 1 subject with 8 sessions and 100 trials per session (from our empirical example, see Section 4) was 01:08 min (ITEM) vs. 16:11 min (LS-S). All computations were performed using SPM12 in MATLAB R2013b running on a 64-bit Windows 7 PC with 16 GB RAM and four hyperthreaded Intel i7 CPU kernels working at 3.40 GHz.

2 This can be seen in equations (9) and (14) by splitting up *U* ^−1^ into 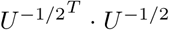, such that the whitening matrix *U* ^−1*/*2^ modifies *T* and 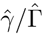 to account for the correlations implied by *U*.

3 This means that trial-wise response amplitudes are only a function of condition-wise response amplitudes, without any further trial-by-trial noise within experimental conditions, i.e. *γ* = *Tβ*.

4 More precisely, the inverse transformed encoding model reads 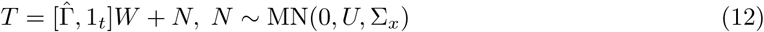 where a *t* × 1 vector of ones is added to the “design matrix” 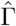 as a constant regressor, such that the model is able to reconstruct discrete differences with arbitrary offsets in the “data matrix” *T*.

5 In this equation, the subscript ¬*j* indicates that 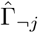 and *T*_*¬j*_ are vertical concatenations and that *U*_*¬j*_ is a block-diagonal combination of the respective session-wise matrices

6 When there are just two classes, *C* can also be a *p* × 1 vector with +1’s and –1’s.

7 Equation (17) uses Iverson bracket notation, i.e. [*p*] = 1, if *p* true and [*p*] = 0, if *p* false.

8 In our implementation of the method (see Section 6), *C* can also be a *p* × 1 vector.

9 Findings denoted as “results not shown” can be found in the sub-folder “Simulation/null results/” of the accompanying GitHub repository (https://github.com/JoramSoch/ITEM-paper)

10 Note that this step would normally not be required when an *a priori* (structural or functional) ROI image is already available or when a searchlight-based ITEM analysis is desired.

11 This means that backwards regression using equations (14) and (15) with *U* = *I*_*t*_ was used for LS-A and LS-S. We also applied support vector regression (SVR) to reconstruct sector intensity levels from LS-A and LS-S estimates, but reconstruction performance was inferior to the results reported here and obtained using ITEM-style inversion (see Section 4.2, Step 7). SVR code is available on the GitHub repository (https://github.com/JoramSoch/ITEM-paper/blob/master/README.md#Application).

12 To be more precise, this requires that (i) inter-stimulus-intervals are sufficiently long (such that HRF regressors do not overlap), (ii) serial correlations are sufficiently weak (such that non-overlapping HRF regressors are not mixed by *V*), (iii) there are no regressors of no interest (such that there are no trial-by-covariate correlations) and (iv) each trial is convolved with the same HRF (such that *U* is not only a diagonal matrix, but a product of the identity matrix).

